# Antigenic evolution of human influenza H3N2 neuraminidase is constrained by charge balancing

**DOI:** 10.1101/2021.07.10.451918

**Authors:** Yiquan Wang, Ruipeng Lei, Armita Nourmohammad, Nicholas C. Wu

## Abstract

As one of the main influenza antigens, neuraminidase (NA) in H3N2 virus has evolved extensively for more than 50 years due to continuous immune pressure. While NA has emerged as an effective vaccine target recently, biophysical constraints on the antigenic evolution of NA remain largely elusive. Here, we apply deep mutational scanning to characterize the local fitness landscape in an antigenic region of NA in six different human H3N2 strains that were isolated around 10 years apart. The local fitness landscape correlates well among strains and the pairwise epistasis is highly conserved. Our analysis further demonstrates that local net charge governs the pairwise epistasis in this antigenic region. In addition, we show that residue coevolution in this antigenic region can be predicted by charge states and pairwise epistasis. Overall, this study demonstrates the importance of quantifying epistasis and the underlying biophysical constraint for building a predictive model of influenza evolution.

## Introduction

There are two major antigens on the surface of influenza virus, hemagglutinin (HA) and neuraminidase (NA). Although influenza vaccine development has traditionally focused on HA, NA has emerged as an effective vaccine target in the past few years, because recent studies have shown that NA immunity has a significant role in protection against influenza infection^1–5^. Influenza NA has an N-terminal transmembrane domain, a stalk domain, and a C-terminal head domain. The head domain of NA functions as an enzyme to cleave the host receptor (i.e. sialylated glycan), which is essential for virus release. Most NA antibodies target the surface loop regions that surround the highly conserved catalytic active site^6–12^. Due to the need to constantly escape from herd immunity (also known as antigenic drift), both HA and NA of human influenza virus have evolved extensively^13–15^. For example, since influenza H3N2 virus entered the human population in 1968, its HA and NA have accumulated more than 83 and 73 amino acid mutations, respectively (Supplementary Fig. 1), which accounted for ∼15% of their protein sequences. However, the evolution of NA is much less well characterized as compared to HA.

To understand how the evolutionary trajectories of NA are being shaped, it is important to characterize the underlying biophysical constraints that govern the fitness of individual amino acid mutations and epistatic interactions between mutations^16–18^. Epistasis is a phenomenon in which the fitness effect of a mutation is dependent on the presence or absence of other mutations. Since epistasis can lead to differential fitness effects of a given mutation on different genetic backgrounds, it can restrict evolutionary trajectories or open up a new functional sequence space that would otherwise be inaccessible^19^. As a result, epistasis is a primary challenge for predicting evolution^20,21^. Nevertheless, epistasis is pervasive in natural evolution in general^22^, and has been shown to influence the evolution of influenza virus^18,19,23–31^. While epistasis is critical for the emergence of oseltamivir-resistant mutants of influenza NA^28–31^, the role of epistasis and the underlying biophysical constraints on NA antigenic evolution remain largely unclear.

Deep mutational scanning combines saturation mutagenesis and next-generation sequencing to determine the phenotypic effects of numerous mutations in a highly parallel manner^32–34^. Deep mutational scanning has been employed to measure the replication fitness effect of all possible single amino acid mutations across different influenza proteins^32,35–38^, which in turn can help to model and predict the natural evolution of human influenza virus^32,35^. More recently, deep mutational scanning is also applied to examine the local fitness landscape and to identify epistasis in influenza HA^24,26,39^. By dissecting the mechanistic basis of epistasis, these studies have provided important insight into the biophysical constraints on HA antigenic evolution^24,26^.

In this study, we aim to understand how epistasis influence NA antigenic evolution and investigate the underlying biophysical constraints. Specifically, we used deep mutation scanning to characterize the local fitness landscape of a NA antigenic region in six different human H3N2 strains that were isolated ∼10 years apart. Our results indicate that the local fitness landscape of this NA antigenic region is highly correlated across six different genetic backgrounds. In-depth analyses further demonstrate that local net charge balancing is a biophysical constraint that governs the epistasis within this NA antigenic region. Lastly, we show that epistasis can help predict residue coevolution in naturally circulating influenza strains.

## Results

### Deep mutational scanning of an antigenic region on NA

When the structure of N2 NA was first reported in 1983^40^, seven regions (I to VII) were proposed to be targeted by antibodies^9^. Within regions I to III, mutations at residues 329, 344, 368, and 370 have been shown to escape monoclonal antibodies^6,41,42^. These four residues, along with residues 328 (region I), 367 (region III), and 369 (region III), form a cluster of seven residues that are very close in space (Fig. 1a-b). Both residues 328 and 367 are under positive selection in human H3N2 NA^43^, whereas coevolution of residues 367 and 369 in human H3N2 NA has created a N-glycosylation site (NXT) at residue 367 during 2010 (Fig. 1c). These observations indicate that residues 328, 367, and 369 also participated in the antigenic drift of human H3N2 NA. By focusing on this seven-residue antigenic region, this study aimed to dissect the biophysical constraints on NA antigenic evolution.

**Figure 1.**
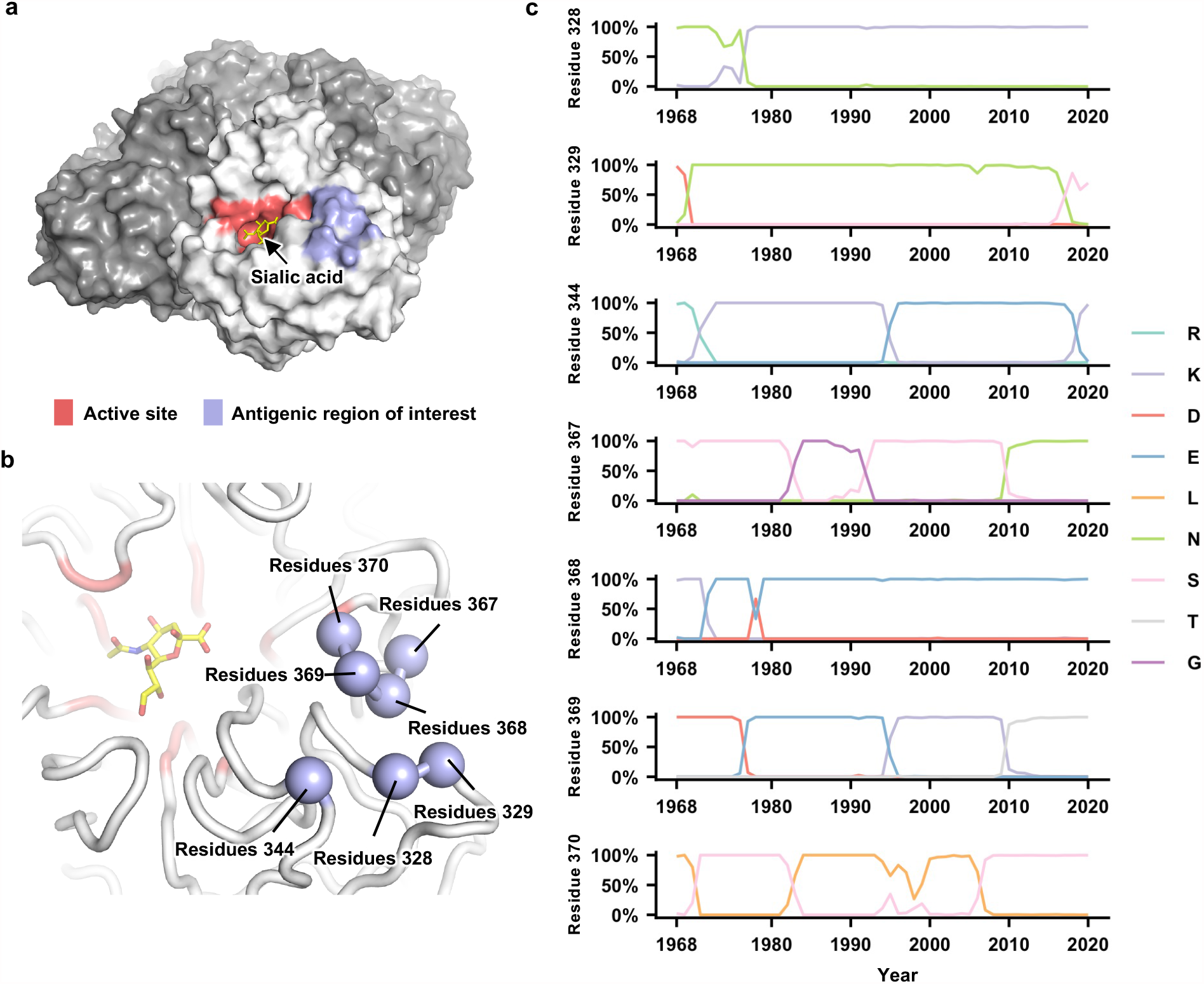
Natural evolution of an antigenic region in human H3N2 NA. **(a)** The enzyme active site and antigenic site are highlighted as red and blue, respectively, on one protomer of the neuraminidase (NA) tetramer. The sialic acid within the active site is shown in yellow. **(b)** Seven residues in the antigenic region of interest are highlighted in blue spheres. **(c)** Natural occurrence frequencies of the amino acid variants that have a natural occurrence at >50% in any given year at the residues of interest are shown. Of note, only those amino acid variants with a natural occurrence at >80% in any given year were included in our mutant libraries. Therefore, although D368 is shown in this plot, it was not included in our mutant libraries since it only reached a maximum occurrence of 67%.

We first compiled a list of amino acid variants at NA residues 328, 329, 344, 367, 368, 369, and 370 that reached an occurrence frequency of >80% in any given year during the natural evolution of human H3N2 viruses since 1968 (Fig. 1c). This list includes two amino acid variants at residue 328 (Asn and Lys), three at residue 329 (Asn, Ser, and Asp), three at residue 344 (Lys, Glu, and Arg), three at residue 367 (Asn, Ser, and Gly), two at residue 368 (Lys and Glu), four at residue 369 (Lys, Glu, Asp, and Thr), and two at residue 370 (Leu and Ser). Together, there are 2 × 3 × 3 × 3 × 2 × 4 × 2 = 864 possible amino acid combinations (also called haplotypes) across these seven residues, although only 53 of them have been observed in naturally circulating human H3N2 strains (Supplementary Fig. 2). Using PCR primers that carry degenerate nucleotides (see Methods and Supplementary Table 1), these 864 variants were introduced into the NA of six different strains (genetic backgrounds) from 1968 to 2019, namely A/Hong Kong/1/1968 (HK68), A/Bangkok/1/1979 (Bk79), A/Beijing/353/1989 (Bei89), A/Moscow/10/1999 (Mos99), A/Victoria/361/2011 (Vic11), and A/Hong Kong/2671/2019 (HK19). All these six strains were historical vaccine strains and isolated approximately 10 years apart.

To measure the virus replication fitness of all 864 variants in the six different genetic backgrounds of interest (i.e. HK68, Bk79, Bei89, Mos99, Vic11, and HK19), deep mutational scanning was employed. Subsequently, six different local fitness landscapes, each with 864 variants, were obtained. Fitness value of each variant was normalized to the corresponding wild type (WT), such that the WT fitness value was 0, whereas positive and negative fitness values represented beneficial and deleterious variants, respectively. Two biological replicates of each experiment were performed. Overall, a high correlation was observed between the replicates (Pearson correlation = 0.81 to 0.89) except for HK68 and Bk79 (0.39 and 0.63, respectively), mostly due to the high measurement noise for low fitness variants (Supplementary Fig. 3).

### Dynamics of the local fitness landscape across genetic backgrounds

To examine whether the local fitness landscape of the NA antigenic region of interest changes over time, fitness measurements of the 864 variants were compared across different genetic backgrounds (Fig. 2a-b). Interestingly, while most variants were strongly deleterious (i.e., had a fitness of <-1) in HK68, variants with a fitness of <-1 were rare in other genetic backgrounds (Fig. 2a). Notably, the difference in variant fitness distribution across genetic backgrounds was not due to the difference in their WT replication fitness since the viral titers from a rescue experiment were similar between HK68 WT, Bei89 WT, and Mos99 WT (Supplementary Fig. 4). In contrast, the fitness of individual variants correlated well across genetic backgrounds (Pearson correlation = 0.48 to 0.79, Fig. 2b), despite their differences in variant absolute fitness values (Fig. 2a). These results demonstrate that although epistasis exists between the seven-residue antigenic region and the rest of the NA sequence, such epistasis is largely variant-nonspecific. Consequently, the topology of the local fitness landscape is highly conserved across genetic backgrounds. This observation is very different from a similar study on a major antigenic site of HA, where the topology of the local fitness landscape differed dramatically among genetic backgrounds^26^.

**Figure 2.**
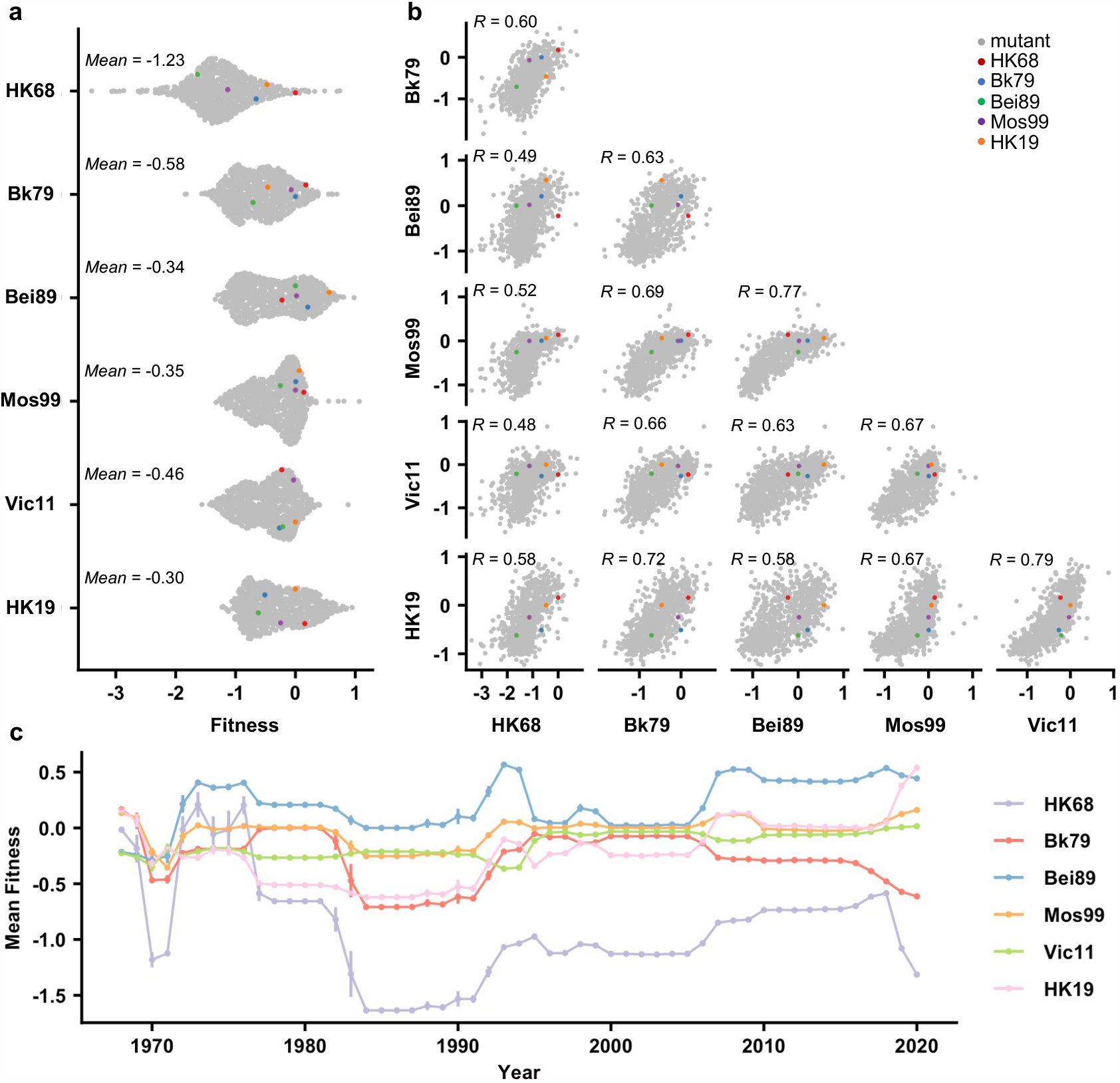
Comparing the local fitness landscapes of the NA antigenic region in six human H3N2 strains. **(a)** Variant fitness distributions in different genetic backgrounds are shown as a sina plot. Each data point represents one variant. A total of 864 data points are present for each row (each genetic background). **(b)** Correlations of variant fitness distribution among different genetic backgrounds are shown, with each data point representing one variant. Pearson correlation coefficients (R) are indicated. **(a, b)** Data points corresponding to the WT sequences of HK68, Bk79, Bei89, Mos99, and HK19 are colored as indicated. Of note, the WT sequence of Vic11 contains a naturally rare variant T329. Therefore, the WT sequence of Vic11 was not included in our mutant libraries. **(c)** Naturally occurring variants were grouped by the year of isolation and their mean fitness in different genetic backgrounds are shown. Different genetic backgrounds are represented by different lines, which are color coded as indicated on the right. Error bars represent the standard error of mean. This analysis included 66,562 NA sequences from human H3N2 strains that were isolated between 1968 and 2020.

We further examined the fitness of naturally occurring variants in different genetic backgrounds (HK68, Bk79, Bei89, Mos99, Vic11, and HK19) (Fig. 2c). Most natural variants have a fitness between -0.5 to 0.5 on the background of Bk79, Bei89, Mos99, Vic11, and HK19. In contrast, most natural variants from 1980s onward are strongly deleterious with fitness <-1 on the background of HK68, indicating many natural variants would not have emerged if the genetic background had not evolved. In other words, the emergence of natural variants in the antigenic region of interest is contingent on the evolution of the rest of the NA sequence. This result is consistent with the observation that HK68 has a much lower mutational tolerance at this antigenic region (Fig. 2a).

### Conservation of pairwise epistasis across genetic backgrounds

To understand the biophysical constraints on NA antigenic evolution, the local fitness landscapes were decomposed into additive fitness effects of individual amino acid variants and pairwise epistasis between amino acid variants^16^. Additive fitness describes the independent contributions of each amino acid variant to fitness, whereas pairwise epistasis describes the non-additive interactions between pairs of amino acid variants. For each genetic background, additive fitness and pairwise epistasis were inferred from the deep mutational scanning data using an established statistical learning model^44^ (see Methods). Model was evaluated using repeated k-fold cross-validation and hyperparameters were chose by maximizing the R^2^ of model prediction and the Pearson correlation coefficient of model parameters (Supplementary Fig. 5). While the correlations between additive fitness contributions varied hugely across genetic backgrounds (Pearson correlation = 0.12 to 0.88, Fig. 3a and c, Supplementary Fig. 6), we observed generally strong correlations between pairwise epistatic effects across the six different genetic backgrounds (Pearson correlation = 0.69 to 0.86, Fig. 3b and c, Supplementary Fig. 7). Overall, these results indicate that pairwise epistasis, but not additive fitness, is highly conserved at the NA antigenic region of interest across genetic backgrounds.

**Figure. 3.**
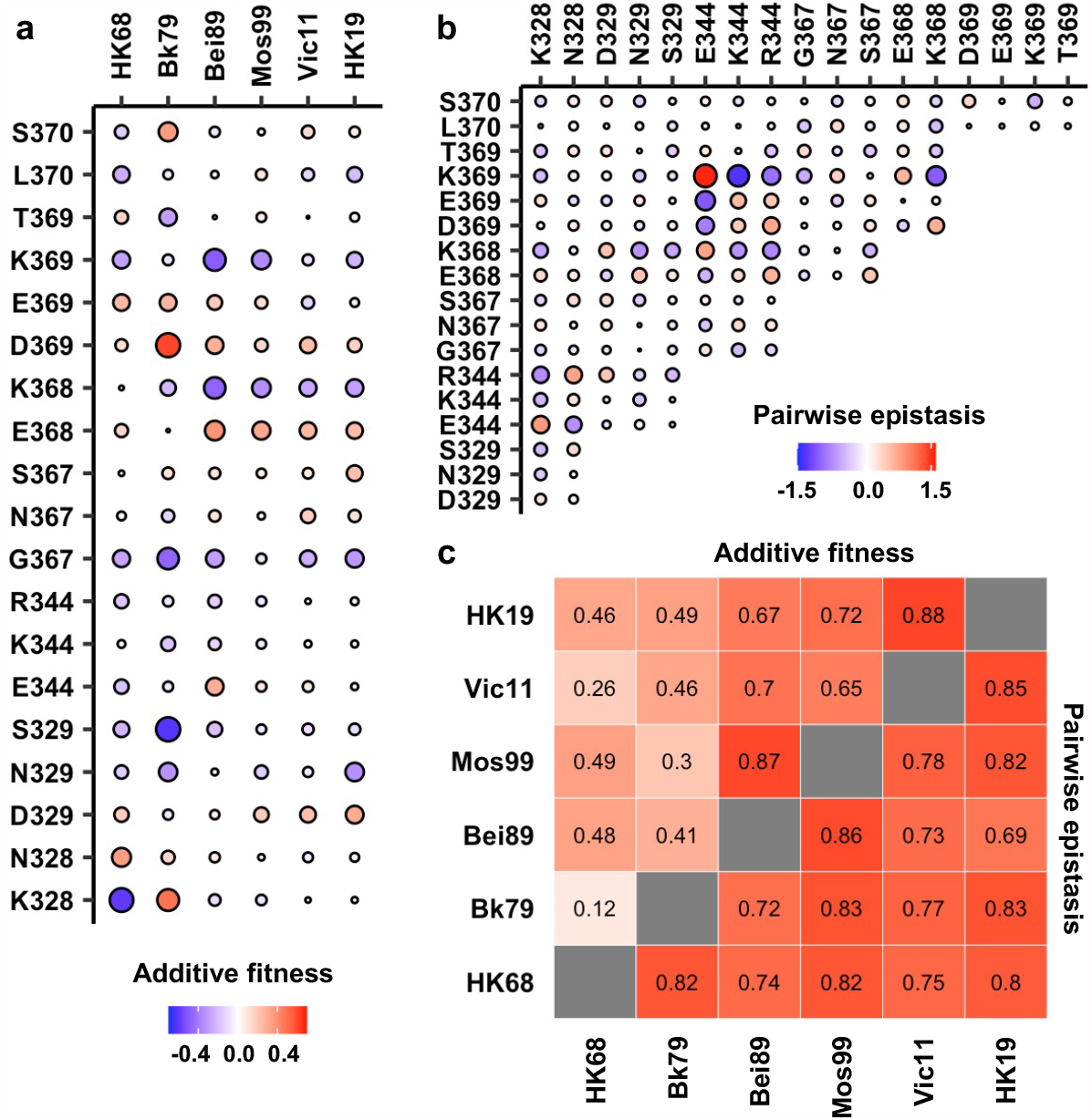
Inference of additive fitness and pairwise epistasis. **(a)** Parameters for additive fitness in different genetic backgrounds are shown. **(b)** Pairwise epistatic effects show strong correlations among six different genetic backgrounds and therefore Bk79 is shown as a representative. The identity of a given double amino acid variant is represented by the labels on the x- and y-axes. The same plots for other genetic backgrounds are shown in Supplementary Fig. 8. **(a, b)** Positive additive fitness and pairwise epistasis are in red. Negative additive fitness and pairwise epistasis are in blue. The magnitude is proportional to the size of the circle. **(c)** Correlation matrices of additive fitness and pairwise epistasis among six genetic backgrounds are shown as a heatmap. See Supplementary Figures 6 and 7 for the related scatter plots.

### Local net charge imposes a biophysical constraint on NA antigenic evolution

Next, we investigated the biophysical constraints that have led to such conserved patterns of epistatic interactions in the NA antigenic region of interest. We noticed that amino acid variants with opposite charges usually exhibited positive epistasis, whereas amino acid variants with the same charge usually exhibited negative epistasis (Fig. 3b and 4a, Supplementary Fig. 8 and 9, see Methods). In contrast, pairwise interactions that involved a neutral amino acid variant did not show any bias towards positive or negative epistasis. These results suggest that balancing of charged amino acid variants imposes a key biophysical constraint on the evolution of this NA antigenic region.

To further probe the mechanism of epistasis in this NA antigenic region, we analyzed a published crystal structure of human H3N2 NA that has K328, E344, and K369 (Fig. 4b)^8^. Both variant pairs E344/K369 and K328/E344 exhibited positive epistasis across all genetic backgrounds, whereas K328/K369 exhibited negative epistasis (Supplementary Fig. 8). In fact, E344 and K369 had the strongest positive epistasis in five of the six genetic backgrounds of interest (except Bei89). Our structural analysis showed that the distance between the side chain carboxylate oxygen of E344 and the side chain amine nitrogen of K369 is 6.6 Å, whereas the distance between the side chain carboxylate oxygen of E344 and the side chain amine nitrogen of K328 is 5.5 Å (Fig. 4b and Supplementary Fig. 10). At these distances, salt bridges cannot be formed and the electrostatic attraction force is negligible^45,46^. Similarly, the distance between the side chain amine nitrogen atoms of K328 and K369 is 11.6 Å (Fig. 4b), which is too far for any significant electrostatic repulsion force^46^. As a result, direct side chain-side chain interaction via electrostatic attraction or repulsion is unlikely to be a major determinant for epistasis in this NA antigenic region. Consistently, pairwise epistasis between two amino acid variants does not correlate with their Cα-Cα distances (Fig. 4d-f and Supplementary Fig. 11), further substantiating that direct side chain-side chain interaction is not a determinant for epistasis here.

**Figure 4.**
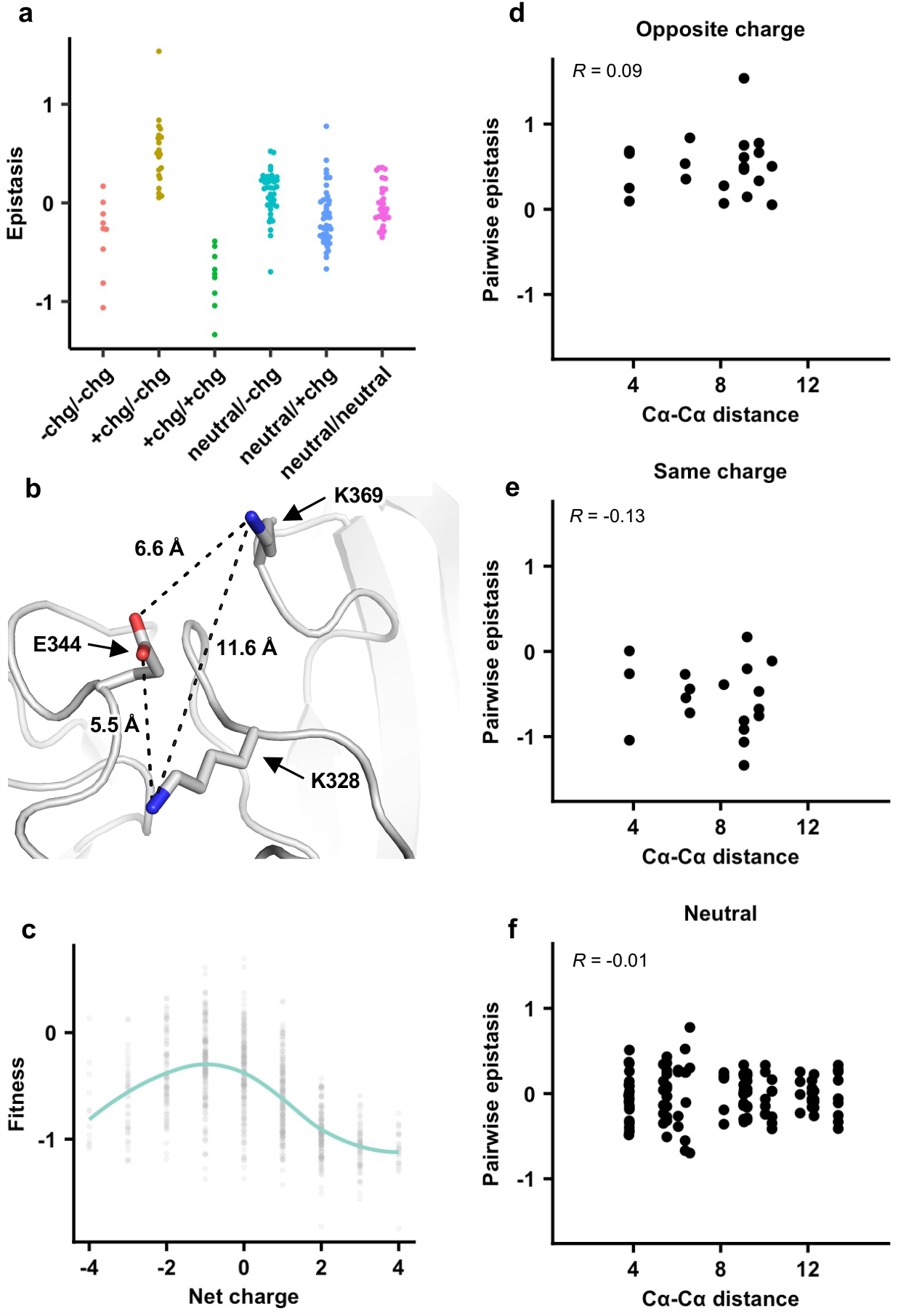
The importance of local net charge in the NA antigenic region. **(a)** Pairwise epistasis in genetic background Bk79 is shown and classified based on amino acid charges. +chg represents positively charged amino acids (K/R), -chg represents negatively charged amino acids (D/E) and neutral represents the remaining amino acids. The same plots for other genetic backgrounds are shown in Supplementary Fig. 9. **(b)** NA structure from H3N2 A/Memphis/31/98 (PDB 2AEP), which has K328, E344, and K369, was analyzed^6^. A similar NA structure from H3N2 A/Tanzania/205/2010, which has K328 and E344, is shown in Supplementary Fig. 10. **(c)** The relationship between variant fitness and net charge in genetic background Bk79 is shown. A smooth curve was fitted by loess and shown in teal. The same plots for other genetic backgrounds are shown in Supplementary Fig. 12. **(d-f)** Relationship between the Cα-Cα distances and epistasis for **(d)** variant pairs with opposite charges, **(e)** variant pairs with the same charge, and **(f)** variant pairs that involve a neutral amino acid in genetic background Bk79 is shown. Pearson correlation coefficient (R) is indicated.

We then analyzed the relationship between variant fitness and local net charge as a molecular phenotype. Here, local net charge was defined as the sum of charges at the seven residues of interest (residues 328, 329, 344, 367, 368, 369, and 370), where positively charged amino acids (K and R) was +1 and negatively charged residues (D and E) was -1. In all genetic backgrounds, variants tended to have a higher fitness when the local net charge was around -1, while variants with a more positive or negative local net charge usually had a lower fitness (Fig. 4c and Supplementary Fig. 12). Overall, our results demonstrate that the local net charge is a key molecular phenotype with a biophysical function that is under balancing selection. This biophysical phenotype imposes a strong constraint on the evolution of the NA antigenic region, reflected in the conserved epistatic interactions between amino acid variants of this region.

### Using local net charge and epistasis to predict NA evolution

Next, we aimed to understand whether the local net charge at the NA antigenic region of interest influences its evolution in circulating human H3N2 strains. We retrieved 66,562 human H3N2 NA sequences spanning from 1968 to 2020 from the Global Initiative for Sharing Avian Influenza Data (GISAID)^47^. Most natural variants in the antigenic region of interest had a local net charge between -1 and +1 (Fig. 5a). Natural variants with a local net charge of -3, -2, +2, or +3 could also be observed but were rare. This observation suggests that the natural evolution of this NA antigenic region is constrained by balancing the local net charge.

**Figure 5.**
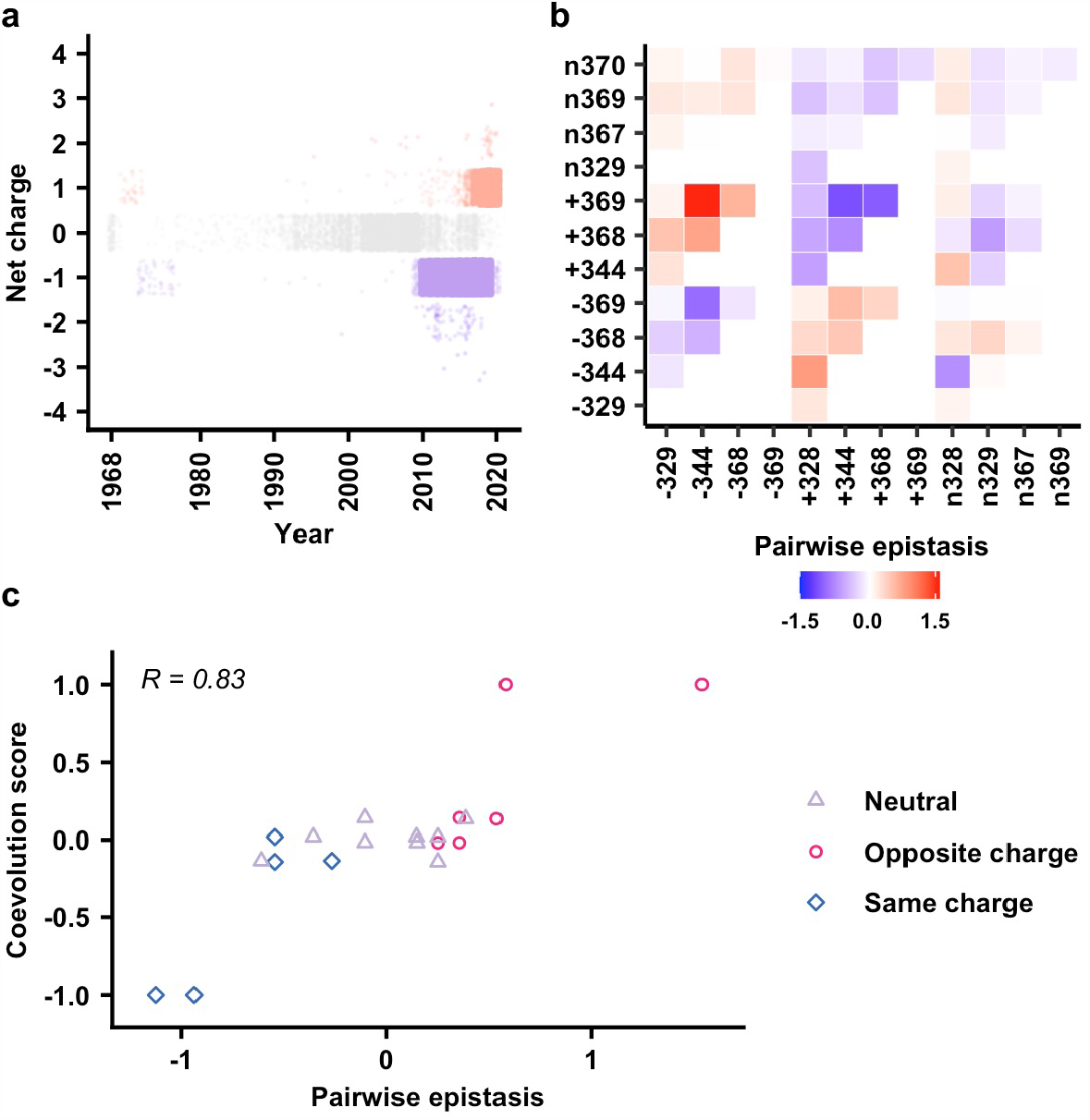
Predicting coevolution of charge states in the NA antigenic region using epistasis. **(a)** The evolution of local net charge at the NA antigenic region of interest is shown. **(b)** Pairwise epistasis of charge states among different residues. Amino acids were classified based on charges. (+) represents positively charged amino acids (K/R/H). (-) represents negatively charged amino acids (D/E). (n) represents the remaining amino acids. **(c)** Relationship between coevolution score and pairwise epistasis in genetic background Bk79 is shown. Pearson correlation coefficient (R) is indicated. The same plots for other genetic backgrounds are shown in Supplementary Fig. 16.

We further tested whether epistasis can be used to predict the evolution of local net charge at the NA antigenic region of interest. A coarse-grained analysis that only considered three charged states, namely positive, negative, and neutral for each residue was performed (see Methods, Supplementary Fig. 13). In this analysis, amino acids were classified into (+) as positively charged (amino acids K/R), (-) as negatively charged (amino acids D/E), and (n) as neutral representing the remaining amino acids. We then computed the pairwise epistasis between different charge classes in two loci by averaging over all the corresponding amino acids in each class. For example, the epistasis between positive charge at residue 344 and negative charge at residue 369 (i.e., +344/-369) is the averaged epistasis over K344/D369, R344/D369, K344/E369, and R344/E369 (Fig. 5b and Supplementary Fig. 14). In addition, we evaluated a coevolution score between a pair of charged states at two different residues (Methods, Supplementary Fig. 15, Supplementary Table 3). In our definition, two charge states that emerged or disappeared shortly one after another would have a positive coevolution score. In contrast emergence of a charge state in one residue follow by disappearance of a charge state in another residue would result in a negative coevolution score. Since residues 367 and 370 were dominated by neutral charge since 1968, they were not included on our coevolution analysis (Supplementary Fig. 13). When we compared the coevolution score with the pairwise epistasis (Fig. 5c and Supplementary Fig. 16), high correlation was observed (Pearson correlation = 0.73 to 0.89). Specifically, pairs with opposite charges usually have positive coevolution scores and positive epistasis, whereas pairs with the same charge usually have negative coevolution scores and negative epistasis. Overall, our analysis demonstrates a biophysically grounded epistatic landscape which can be used to predict the natural coevolution of residues in this NA antigenic region.

## Discussion

Understanding the biophysical constraints on evolution is a key to building a predictive evolutionary model^17,21,48–52^ Through a systematic analysis of pairwise epistasis, this study shows that the local net charge is a major biophysical molecular phenotype that constrains the evolution of an antigenic region in human influenza H3N2 NA. An important feature for this antigenic region of interest is that the pairwise epistasis are highly conserved across diverse genetic backgrounds. The conservation of pairwise epistasis and the constraint on local net charge subsequently enable a predictive modelling of residue coevolution in this NA antigenic region in naturally circulating human influenza H3N2 virus.

A key finding of this study is that the optimal local net charge at the antigenic region of interest is slightly negative, and an increase or decrease in local net charge is deleterious. This characteristic suggests that the local net charge may be the key molecular phenotype under stabilizing and balancing selection, imposing a biophysical constraint on evolution of the NA antigenic region. This local net charge may have several non-exclusive functional roles. First, since the host cell membrane and sialylated glycan receptor are both negatively charged, charge distribution on the virus surface, including the antigenic region of interest, may affect the kinetics of host membrane attachment and virus release. Second, the antigenic site of interest is proximal to the catalytic active site (Fig. 1a), the local net charge may affect the catalytic efficiency of NA. Lastly, the local net charge may influence the protein stability of NA^53,54^. Understanding the detailed molecular mechanisms of biophysical constraints, albeit beyond the scoop of this study, will likely further enhance the ability to predict evolution.

One interesting observation in this study is that the ancestral strain HK68 has a much lower mutational tolerability at the antigenic region of interest than the subsequent strains, although the topology of the local fitness landscapes is largely conserved across strains. This result suggests that other biophysical features of NA are evolving over time and that they epistatically interact with the antigenic region of interest in a residue nonspecific manner. Nonspecific epistasis is often related to protein stability^16^, which is best described by the threshold robustness model^55^. Under the threshold robustness model, slightly destabilizing mutations may have neutral fitness when the protein has an excess stability margin but are deleterious when the protein is marginally stable. Threshold robustness model can be used to explain the differential variant fitness distribution among the six genetic backgrounds in our study. For example, it is possible that Mos99 NA has an excess stability margin such that many variants are nearly neutral despite being slightly destabilizing. In contrast, the same slightly destabilizing variants are highly deleterious in HK68 NA because it is marginally stable. When comparing the variant fitness in different genetic backgrounds, some nonlinearity can be observed (Fig. 2b), which is a feature of the threshold robustness model^49,56^. Nevertheless, additional studies are needed to confirm whether the difference in variant fitness distribution among genetic backgrounds is due to protein stability or other biophysical factors.

Predicting the evolution of human influenza virus is a challenging task, yet important for seasonal influenza vaccine development. Constructing a predictive model of human influenza evolution has shown to benefit from phylogenetic information^21^, antigenic data^57^, in vitro measurements of mutant fitness effects^32^, and experimental selection for antibody escape variants^58^. This work further illustrates that a biophysical epistatic model of antigenic fitness landscape can also be instrumental in predicting the evolution of human influenza virus. As the knowledge about the evolutionary biology of influenza virus accumulates, a unifying model that can accurately predict emerging mutation may one day be built.

## Methods

### Recombinant influenza virus

All H3N2 viruses generated in this study were based on the influenza A/WSN/33 (H1N1) eight-plasmid reverse genetics system^59^. Chimeric 6:2 reassortments were employed with the HA ectodomains from the H3N2 A/Hong Kong/1/1968 (HK68) and the entire neuraminidase (NA) coding region from the strains of interest^39^. For HA, the ectodomain was from HK68, whereas the non-coding region, N-terminal secretion signal, C-terminal transmembrane domain, and cytoplasmic tail were from A/WSN/33. For NA, the entire coding region was from the strains of interest, whereas the non-coding region of NA was from A/WSN/33. H3N2 strains of interest in this study were as follows with Global Initiative for Sharing Avian Influenza Data (GISAID)^47^ accession numbers in parentheses: A/Hong Kong/1/1968 (EPI_ISL_245769), A/Bangkok/1/1979 (EPI_ISL_122020), A/Beijing/353/1989 (EPI_ISL_123212), A/Moscow/10/1999 (EPI_ISL_127595), A/Victoria/361/2011 (EPI_ISL_158723), and A/Hong Kong/2671/2019 (EPI_ISL_882915).

### Mutant library construction

For each mutant library, insert and vector fragments were generated by PCR using PrimeSTAR Max DNA Polymerase (Takara) according to the manufacturer’s instructions, with wild-type (WT) NA-encoding plasmid (pHW2000-NA) as templates. Primers for the insert contained the combinatorial mutations of interest and are shown in Supplementary Table 1. For insert fragment, two rounds of PCR were performed. Forward primers (P2 set) and reverse primers (P3 set) were mixed at the indicated molar ratio and used for the first round PCR (Supplementary Table 1). Products from the first round PCR were then purified using Monarch DNA Gel Extraction Kit (New England Biolabs) and used as the templates for the second round insert PCR. Forward primers (P1 set) and reverse primers (P3 set) were mixed at the indicated molar ratio and used for the second round PCR (Supplementary Table 1). Of note, the same reverse primers (P3 set) were used for both rounds PCR. Primers for the vector PCR are also shown in Supplementary Table 1. The final PCR products of the inserts and vectors were purified by PureLink PCR purification kit (Thermo Fisher Scientific), digested by DpnI and BsmBI (New England Biolabs), and ligated using T4 DNA ligase (New England Biolabs). The ligated product was transformed into MegaX DH10B T1R cells (Thermo Fisher Scientific). At least one million colonies were collected for each mutant library. Plasmid mutant libraries were purified from the bacteria colonies using Plasmid Midi Kit (Qiagen).

### Deep mutational scanning

Each plasmid mutant library was transfected into HEK293T/MDCK-SIAT1 cells co-culture (ratio of 6:1) at 60% confluence in a T75 flask, using lipofectamine 2000 (Thermo Fisher Scientific) according to the manufacturer’s instructions. At 24 hours post-transfection, cells were washed twice with phosphate-buffered saline (PBS) and cell culture medium was replaced with OPTI-MEM medium containing 0.8 µg mL^-1^ TPCK-trypsin. After 72 hours post-transfection, virus was harvested and titered by TCID50 assay using MDCK-SIAT1 cells then stored at -80°C until use. To passage the virus mutant libraries, MDCK-SIAT1 cells in s T75 flask were washed twice with PBS and then infected with a multiplicity of infection (MOI) of 0.02 in OPTI-MEM medium containing 0.8 µg mL^-1^ TPCK-trypsin. Infected cells were washed twice with PBS at 2 hours post-infection then fresh OPTI-MEM medium containing 0.8 µg mL^-1^ TPCK-trypsin was added to the cells. At 24 hours post-infection, supernatant containing the virus was collected. Viral RNA was extracted using QIAamp Viral RNA Mini Kit (Qiagen). Purified viral RNA was reverse transcribed to cDNA using Superscript III reverse transcriptase (Thermo Fisher Scientific). The adapter sequence for Illumina sequencing was added to the plasmid or cDNA from the post-passaging virus mutant libraries by PCR using sequencing library preparation primers listed in Supplementary Table 1. An additional PCR was performed to add the rest of the adapter sequence and index to the amplicon using primers: 5’-AAT GAT ACG GCG ACC ACC GAG ATC TAC ACT CTT TCC CTA CAC GAC GCT-3’ and 5’-CAA GCA GAA GAC GGC ATA CGA GAT XXX XXX GTG ACT GGA GTT CAG ACG TGT GCT-3’. Positions annotated by an X represented the nucleotides for the index sequence. The final PCR products were purified by PureLink PCR purification kit (Thermo Fisher Scientific) and submitted for next-generation sequencing using Illumina MiSeq PE250.

### Sequencing data analysis

Sequencing data was obtained in FASTQ format and were analyzed using custom python scripts. Briefly, sequences were parsed by SeqIO module in BioPython^60^. After trimming the primer sequences, both forward and reverse-complement of the reverse reads were translated into protein sequences. A paired-end read was then filtered and removed if the protein sequences of the forward and reverse-complement of the reverse reads did not match. Subsequently, amino acids at the residues of interest were extracted. The number of reads corresponding to each of the 864 variants was then counted. The unnormalized fitness of each variant *i* in each replicate was estimated as follow:

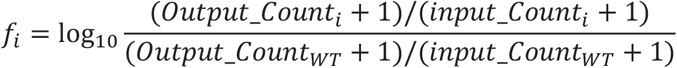

where the *Output_Count*_*i*_ represents the number of reads corresponding to variant *i* in the post-passaging virus mutant library, and the *input_Count*_*i*_ represents the number of reads corresponding to variant *i* in the plasmid mutant library. A pseudocount of 1 was added to the counts to avoid division by zero. Of note, the WT sequence of Vic11 contains a naturally rare variant T329. As a result, the WT sequence of Vic11 was not included in our mutant library design. However, due to incomplete DpnI digestion of the vector during mutant library construction, the WT sequence of Vic11 was present in the Vic11 mutant library and could be detected in the next-generation sequencing data.

The final fitness value for each mutant is:

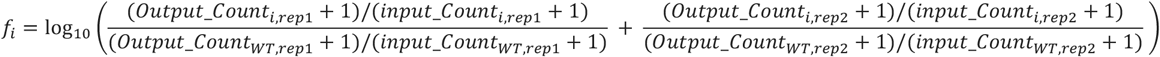

where *rep1* and *rep2* represent replicate 1 and replicate 2, respectively.

### Total charge of residues of interest

Positively charged amino acids (K/R) were assigned with a charge of 1, negatively charged amino acids (D/E) were -1, and neutral amino acids were 0. The net charge of a given variant is defined as the algebraic sum of the charges of the seven residues of interest. For example, the net charge of KNEGKKL, which has three positively charged amino acids and one negatively charged amino acid, is 2.

### Modeling fitness and decomposition of interactions

We modeled variant fitness as a nonlinear function of the sum of the additive and the pairwise effects by MAVE-NN^44^. Statistical learning model was trained to estimate the latent phenotype *ϕ* and prediction fitness *ŷ*.

The sum of the additive and the pairwise effects was defined as a latent phenotype *ϕ*_*pairwise*_,

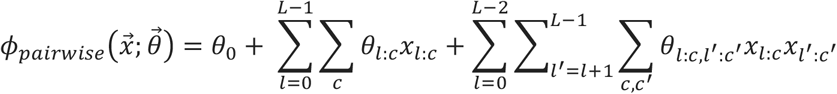

where *L* is the length of the sequences, *C* is the total number of amino acids, and 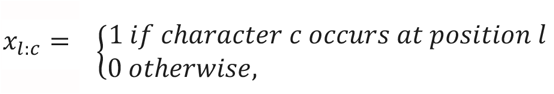 is the one-hot encoding of the sequence at position *l* when amino acid is *c*. 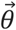 represents the weight of the additive and the pairwise effects. Fitness is then modeled as a nonlinear function of the sum of tanh,

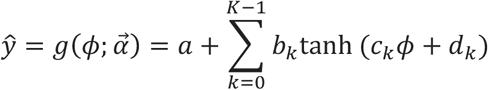

where *K* specifies the number of “hidden nodes” contributing to the sum, and 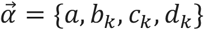 are the trainable parameters.

To train the above model, dataset was reformatted to a set of *N* observations, 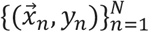, where each observation comprises of sequence 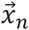 and its fitness *y*_*n*_. Then, the dataset was randomly divided into a training set, a validation set, and a test set with a ratio of 0.64: 0.16: 0.2. Model was evaluated using repeated k-fold cross-validation and hyperparameters were chose by maximizing the R^2^ of model prediction and Pearson correlation coefficient of model parameters^61^. Notably, we find that the R^2^ of our model prediction is insensitive to regularization, and largely depends on the quality of the deep mutational scanning data (Supplementary Fig. 3).

### Estimating epistasis between charged states

Amino acids are classified according to charges. Positive (+) represents positively charged amino acids (K/R), negative (-) represents negatively charged amino acids (D/E), and neutral (n) represents the remaining amino acids. All variants of the NA antigen region are then converted into charge states, named positive, negative, and neutral for each residue. For example, -344 represents K344 and R344. The epistasis value of a given charge state is the average over the epistasis values of all amino acid variants in the specified charge state.

### Analysis of natural sequences

A total of 66,562 full-length NA protein sequences from human H3N2 were downloaded from the GISAID (http://gisaid.org)^47^ (Supplementary Table 2). Amino acid sequences of NA residues 328, 329, 344, 367, 368, 369, and 370 in individual strains were extracted. Individual sequences were grouped by the year of isolation and their mean fitness in different genetic backgrounds were plotted in Fig. 2c. The human H3N2 NA protein sequences used in this study are listed in Supplementary Data 1.

### Inference of natural coevolution score

The change in the natural frequency of the charge states (+/-/n) at residue *i* from year *n* – 1 to year *n* was computed as:

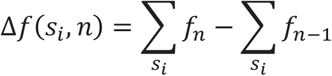

where *s*_*i*_ is the change in the natural frequency of the charge states (+/-/n) at residue *i*, and *f*_*n*_ is the natural frequency of the amino acid in year *n*.

All local maxima were then selected and defined as peak of frequency change using find_peaks module in SciPy. Subsequently, to evaluate the proximity of peaks among two charge states, a coevolution score was calculated as the sum of exponential weights of all pairwise peak distances from a given mutation pair (*i, j*):

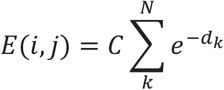

where *d*_*k*_ is the time separation (in years) between two peaks and *C* is initial value. *C* = 1 if both peaks have the same sign, otherwise *C* = -1 (i.e. one peak is positive and the other is negative). See Supplementary Fig. 15 for a schematic overview of calculating the coevolution score.

## Data availability

Raw sequencing data have been submitted to the NIH Short Read Archive under accession number: BioProject PRJNA742436.

## Code availability

Custom python scripts for analyzing the deep mutational scanning data have been deposited to https://github.com/Wangyiquan95/NA_EPI.

## Funding acknowledgement

This work was supported by DFG grant (SFB1310) for Predictability in Evolution (A.N.), MPRG funding through the Max Planck Society (A.N.), the Royalty Research Fund from the University of Washington (A.N.), and National Institutes of Health (NIH) R00 AI139445 (N.C.W.).

## Author Contributions

Y.W. and N.C.W. conceived and designed the study. Y.W. and R.L. performed the deep mutational scanning experiments. Y.W, A.N. and N.C.W. analyzed the data. Y.W., R.L. and N.C.W. wrote the paper and all authors reviewed and/or edited the paper.

## Competing Interests

The authors declare no competing interests.

## Notes

https://www.ncbi.nlm.nih.gov/bioproject/?term=PRJNA742436

### Competing Interest Statement

The authors have declared no competing interest.

