## Supplementary Tables and Figures for "Antigenic evolution of human influenza H3N2 neuraminidase is constrained by charge balancing"

### Supplementary Table 1. Primers for deep mutational scanning experiments.

| Strain | PCR reaction | Primer name | Molar ratio | Sequence (from 5' to 3') |
|---|---|---|---|---|
| HK68 | Mutant library insert | NA_hk68_P1-1F | 2 | ACGCCGTCTCACCTAGAAASRACGACAGATCTAGCAATAGCAATTGCAGGAATCCTAAT |
| | | NA_hk68_P1-2F | 1 | ACGCCGTCTCACCTAGAAASAGCGACAGATCTAGCAATAGCAATTGCAGGAATCCTAAT |
| | | NA_hk68_P2-1F | 2 | GCAATAGCAATTGCAGGAATCCTAATAATGAGARAGGGAATCAAGGAGTGAAAGG |
| | | NA_hk68_P2-2F | 1 | GCAATAGCAATTGCAGGAATCCTAATAATGAGGAAGGGAATCAAGGAGTGAAAGG |
| | | NA_hk68_P3-1R | 2 | AGTACGTCTCATAACCTGAGCGTRAMTCCTYGYTGATCGTTCTTCCCATCCACA |
| | | NA_hk68_P3-2R | 2 | AGTACGTCTCATAACCTGAGCGTRATKTCTYGYTGATCGTTCTTCCCATCCACA |
| | | NA_hk68_P3-3R | 1 | AGTACGTCTCATAACCTGAGCGTRATKTCTYGCCGATCGTTCTTCCCATCCACA |
| | | NA_hk68_P3-4R | 1 | AGTACGTCTCATAACCTGAGCGTRAMTCCTYGCCGATCGTTCTTCCCATCCACA |
| | Mutant library vector | NA_hk68_VF | 1 | AGCTCGTCTCGGTTATGAAACTTTCAAAGTCATTGGTGG |
| | | NA_hk68_VR | 1 | CGACCGTCTCCTAGGTGTGTCGCCAACAAGCCCTGAGCA |
| | Sequencing library preparation | HK68_7mutlib_recover-F | 1 | CACTCTTTCCCTACACGACGCTCTTCCGATCTTGCTCAGGGCTTGTTGGCGACACA |
| | | HK68_7mutlib_recover-R | 1 | GACTGGAGTTCAGACGTGTGCTCTTCCGATCTAATGACTTTGAAAGTTTCATAACC |
| Bk79 | Mutant library insert | NA_bk79_P1-1F | 2 | ACGCCGTCTCACCCAGAAASRACGACAGATCTAGCAGTAGCTATTGCCGGAATCCTAAC |
| | | NA_bk79_P1-2F | 1 | ACGCCGTCTCACCCAGAAASAGCGACAGATCTAGCAGTAGCTATTGCCGGAATCCTAAC |
| | | NA_bk79_P2-1F | 2 | GCAGTAGCTATTGCCGGAATCCTAACAATGAGARAGGGAATCATGGAGTGAAAGG |
| | | NA_bk79_P2-2F | 1 | GCAGTAGCTATTGCCGGAATCCTAACAATGAGGAAGGGAATCATGGAGTGAAAGG |
| | | NA_bk79_P3-1R | 2 | AGTACGTCTCATAACCTGATCGTRAMTCCTYGYTGATCGTTCTTCCCATCCACA |
| | | NA_bk79_P3-2R | 2 | AGTACGTCTCATAACCTGATCGTRATKTCTYGYTGATCGTTCTTCCCATCCACA |
| | | NA_bk79_P3-3R | 1 | AGTACGTCTCATAACCTGATCGTRATKTCTYGCCGATCGTTCTTCCCATCCACA |
| | | NA_bk79_P3-4R | 1 | AGTACGTCTCATAACCTGATCGTRAMTCCTYGCCGATCGTTCTTCCCATCCACA |
| | Mutant library vector | NA_bk79_VF | 1 | AGCTCGTCTCGGTTATGAAACCTTCAAAGTCATTGGTGG |
| | | NA_bk79_VR | 1 | CGACCGTCTCCTGGGTGTGTCGCCAACAAGCCCTGAGCA |
| | Sequencing library preparation | Bk79_7mutlib_recover-F | 1 | CACTCTTTCCCTACACGACGCTCTTCCGATCTTGCTCAGGGCTTGTTGGAGACACA |
| | | Bk79_7mutlib_recover-R | 1 | GACTGGAGTTCAGACGTGTGCTCTTCCGATCTAATGACTTTGAAGGTTTCATAACC |
| Bei89 | Mutant library insert | NA_bei89_P1-1F | 2 | ACGCCGTCTCACCCAGAAASRACGACAGCTCCAGCAGTAGCTATTGCCGGAATCCTAAC |
| | | NA_bei89_P1-2F | 1 | ACGCCGTCTCACCCAGAAASAGCGACAGCTCCAGCAGTAGCTATTGCCGGAATCCTAAC |
| | | NA_bei89_P2-1F | 2 | GCAGTAGCTATTGCCGGAATCCTAACAATGAGARAGGGAGTCATGGAGTGAAAGG |
| | | NA_bei89_P2-2F | 1 | GCAGTAGCTATTGCCGGAATCCTAACAATGAGGAAGGGAGTCATGGAGTGAAAGG |
| | | NA_bei89_P3-1R | 2 | AGTACGTCTCATAACCTGAGCGTRAMTCCTYGYTGATCGTTCTTCCCATCCACA |
| | | NA_bei89_P3-2R | 2 | AGTACGTCTCATAACCTGAGCGTRATKTCTYGYTGATCGTTCTTCCCATCCACA |
| | | NA_bei89_P3-3R | 1 | AGTACGTCTCATAACCTGAGCGTRATKTCTYGCCGATCGTTCTTCCCATCCACA |
| | | NA_bei89_P3-4R | 1 | AGTACGTCTCATAACCTGAGCGTRAMTCCTYGCCGATCGTTCTTCCCATCCACA |
| | Mutant library vector | NA_bei89_VF | 1 | AGCTCGTCTCGGTTATGAAACCTTCAAAGTCATTGGAGG |
| | | NA_bei89_VR | 1 | CGACCGTCTCCTGGGTGTGTCTCCAACAAGCCCTGAGCA |
| | Sequencing library preparation | Bei89_7mutlib_recover-F | 1 | CACTCTTTCCCTACACGACGCTCTTCCGATCTTGCTCAGGGCTTGTTGGAGACACA |
| | | Bei89_7mutlib_recover-R | 1 | GACTGGAGTTCAGACGTGTGCTCTTCCGATCTAATGACTTTGAAGGTTTCATAACC |

| | | | | |
|---|---|---|---|---|
| **Mos99** | **Mutant library insert** | NA_mos99_P1-1F | 2 | ACGCCGTCTCACCCAGAAASRACGACAGCTCCAGCAGTAGCCATTGCTTGGATCCTAAC |
| | | NA_mos99_P1-2F | 1 | ACGCCGTCTCACCCAGAAASAGCGACAGCTCCAGCAGTAGCCATTGCTTGGATCCTAAC |
| | | NA_mos99_P2-1F | 2 | GCAGTAGCCATTGCTTGGATCCTAACAATGAGARAGGTGGTCATGGAGTGAAAGG |
| | | NA_mos99_P2-2F | 1 | GCAGTAGCCATTGCTTGGATCCTAACAATGAGGAAGGTGGTCATGGAGTGAAAGG |
| | | NA_mos99_P3-1R | 2 | AGTACGTCTCATATCCTGAGCGTRAMTCCTYGYTGATCGTTCTTCCCATCCACA |
| | | NA_mos99_P3-2R | 2 | AGTACGTCTCATATCCTGAGCGTRATKTCTYGYTGATCGTTCTTCCCATCCACA |
| | | NA_mos99_P3-3R | 1 | AGTACGTCTCATATCCTGAGCGTRATKTCTYGCCGATCGTTCTTCCCATCCACA |
| | | NA_mos99_P3-4R | 1 | AGTACGTCTCATATCCTGAGCGTRAMTCCTYGCCGATCGTTCTTCCCATCCACA |
| | **Mutant library vector** | NA_mos99_VF | 1 | AGCTCGTCTCGGATATGAAACCTTCAAAGTCATTGAAGG |
| | | NA_mos99_VR | 1 | CGACCGTCTCCTGGGTGTGTCTCCAACAAGTCCTGAGCA |
| | **Sequencing library preparation** | Mos99_7mutlib_recover-F | 1 | CACTCTTTCCCTACACGACGCTCTTCCGATCTTGCTCAGGACTTGTTGGAGACACA |
| | | Mos99_7mutlib_recover-R | 1 | GACTGGAGTTCAGACGTGTGCTCTTCCGATCTAATGACTTTGAAGGTTTCATATCC |
| **Vic11** | **Mutant library insert** | NA_vic11_P1-1F | 2 | ACGCCGTCTCACCCAGAAASRACGACAGCTCCAGCAGTAGCCATTGTTTGGATCCTAAC |
| | | NA_vic11_P1-2F | 1 | ACGCCGTCTCACCCAGAAASAGCGACAGCTCCAGCAGTAGCCATTGTTTGGATCCTAAC |
| | | NA_vic11_P2-1F | 2 | GCAGTAGCCATTGTTTGGATCCTAACAATGAAARAGGTGGTCATGGAGTGAAAGG |
| | | NA_vic11_P2-2F | 1 | GCAGTAGCCATTGTTTGGATCCTAACAATGAAGAAGGTGGTCATGGAGTGAAAGG |
| | | NA_vic11_P3-1R | 2 | AGTACGTCTCATACCCTAAGCGTRAMTCCTYGYTGATTGTTCTTCCCATCCACA |
| | | NA_vic11_P3-2R | 2 | AGTACGTCTCATACCCTAAGCGTRATKTCTYGYTGATTGTTCTTCCCATCCACA |
| | | NA_vic11_P3-3R | 1 | AGTACGTCTCATACCCTAAGCGTRATKTCTYGCCGATTGTTCTTCCCATCCACA |
| | | NA_vic11_P3-4R | 1 | AGTACGTCTCATACCCTAAGCGTRAMTCCTYGCCGATTGTTCTTCCCATCCACA |
| | **Mutant library vector** | NA_vic11_VF | 1 | AGCTCGTCTCGGGTATGAAACCTTCAAAGTCATTGAAGG |
| | | NA_vic11_VR | 1 | CGACCGTCTCCTGGGTGTGTCTCCAACAAGTCCTGAACA |
| | **Sequencing library preparation** | Vic11_7mutlib_recover-F | 1 | CACTCTTTCCCTACACGACGCTCTTCCGATCTTGTTCAGGACTTGTTGGAGACACA |
| | | Vic11_7mutlib_recover-R | 1 | GACTGGAGTTCAGACGTGTGCTCTTCCGATCTAATGACTTTGAAGGTTTCATACCC |
| **HK19** | **Mutant library insert** | NA_hk19_P1-1F | 2 | ACGCCGTCTCACCCAGAAASRACGACAGCTCCAGCAGTAGCCATTGTTTGAATCCTAAC |
| | | NA_hk19_P1-2F | 1 | ACGCCGTCTCACCCAGAAASAGCGACAGCTCCAGCAGTAGCCATTGTTTGAATCCTAAC |
| | | NA_hk19_P2-1F | 2 | GCAGTAGCCATTGTTTGAATCCTAACAATGAAARAGGTGGTCATGGAGTGAAAGG |
| | | NA_hk19_P2-2F | 1 | GCAGTAGCCATTGTTTGAATCCTAACAATGAAGAAGGTGGTCATGGAGTGAAAGG |
| | | NA_hk19_P3-1R | 2 | AGTACGTCTCATACCCTAAGCGTRAMTCCTYGYTGATTGTTCTCCCCATCCACA |
| | | NA_hk19_P3-2R | 2 | AGTACGTCTCATACCCTAAGCGTRATKTCTYGYTGATTGTTCTCCCCATCCACA |
| | | NA_hk19_P3-3R | 1 | AGTACGTCTCATACCCTAAGCGTRATKTCTYGCCGATTGTTCTCCCCATCCACA |
| | | NA_hk19_P3-4R | 1 | AGTACGTCTCATACCCTAAGCGTRAMTCCTYGCCGATTGTTCTCCCCATCCACA |
| | **Mutant library vector** | NA_hk19_VF | 1 | AGCTCGTCTCGGGTATGAAACCTTCAAAGTCGTTGAAGG |
| | | NA_hk19_VR | 1 | CGACCGTCTCCTGGGTGTGTCTCCAACAAGCCCTGAACA |
| | **Sequencing library preparation** | HK19_7mutlib_recover-F | 1 | CACTCTTTCCCTACACGACGCTCTTCCGATCTTGTTCAGGGCTTGTTGGAGACACA |
| | | HK19_7mutlib_recover-R | 1 | GACTGGAGTTCAGACGTGTGCTCTTCCGATCTAACGACTTTGAAGGTTTCATACCC |

### Supplementary Table 2. Human H3N2 NA sequences.

| Year | Number of sequences |
|---|---|
| 1968 | 99 |
| 1969 | 12 |
| 1970 | 10 |
| 1971 | 16 |
| 1972 | 25 |
| 1973 | 10 |
| 1974 | 12 |
| 1975 | 10 |
| 1976 | 17 |
| 1977 | 14 |
| 1978 | 6 |
| 1979 | 4 |
| 1980 | 9 |
| 1981 | 3 |
| 1982 | 12 |
| 1983 | 6 |
| 1984 | 4 |
| 1985 | 12 |
| 1986 | 10 |
| 1987 | 6 |
| 1988 | 14 |
| 1989 | 22 |
| 1990 | 11 |
| 1991 | 40 |
| 1992 | 34 |
| 1993 | 105 |
| 1994 | 75 |
| 1995 | 78 |
| 1996 | 89 |
| 1997 | 97 |
| 1998 | 120 |
| 1999 | 176 |
| 2000 | 222 |
| 2001 | 89 |
| 2002 | 260 |
| 2003 | 486 |
| 2004 | 398 |
| 2005 | 432 |
| 2006 | 344 |
| 2007 | 612 |
| 2008 | 827 |
| 2009 | 1272 |
| 2010 | 1298 |
| 2011 | 1915 |
| 2012 | 2794 |
| 2013 | 2195 |
| 2014 | 4054 |
| 2015 | 5280 |
| 2016 | 6491 |
| 2017 | 11886 |
| 2018 | 8399 |
| 2019 | 13754 |
| 2020 | 2396 |

### Supplementary Table 3. Coevolution score of charged states at different residues

| Charge state i | Charge state j | Coevolution score |
|---|---|---|
| 328 | n329 | 0.019 |
| n328 | -368 | 0.018 |
| n328 | 368 | 0.14 |
| 344 | -369 | 1 |
| n328 | -329 | 0.019 |
| -344 | 369 | 1.00012 |
| 328 | -368 | 0.14 |
| 328 | 368 | 0.018 |
| n329 | -368 | 0.14 |
| -329 | 368 | 0.14 |
| 328 | -329 | -0.019 |
| n328 | -368 | -0.14 |
| n328 | 368 | -0.018 |
| n329 | 368 | -0.14 |
| 328 | -368 | -0.018 |
| 344 | 369 | -1.00012 |
| n328 | n329 | -0.019 |
| -344 | -369 | -1 |
| 328 | 368 | -0.14 |
| -329 | -368 | -0.14 |

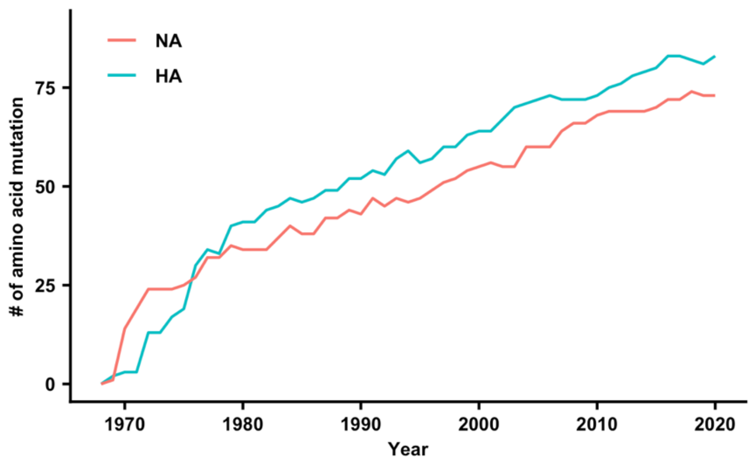


**Supplementary Figure 1. Accumulation of amino acid mutations in the hemagglutinin (HA) and neuraminidase (NA) of human H3N2 viruses.** The number of amino acid mutations in the HA and NA relative to the ancestral strain, A/Hong Kong/1/1968 (HK68) is shown as a line graph. Consensus sequence of all isolates from a given year is used for calculating the number of mutations relative to HK68.

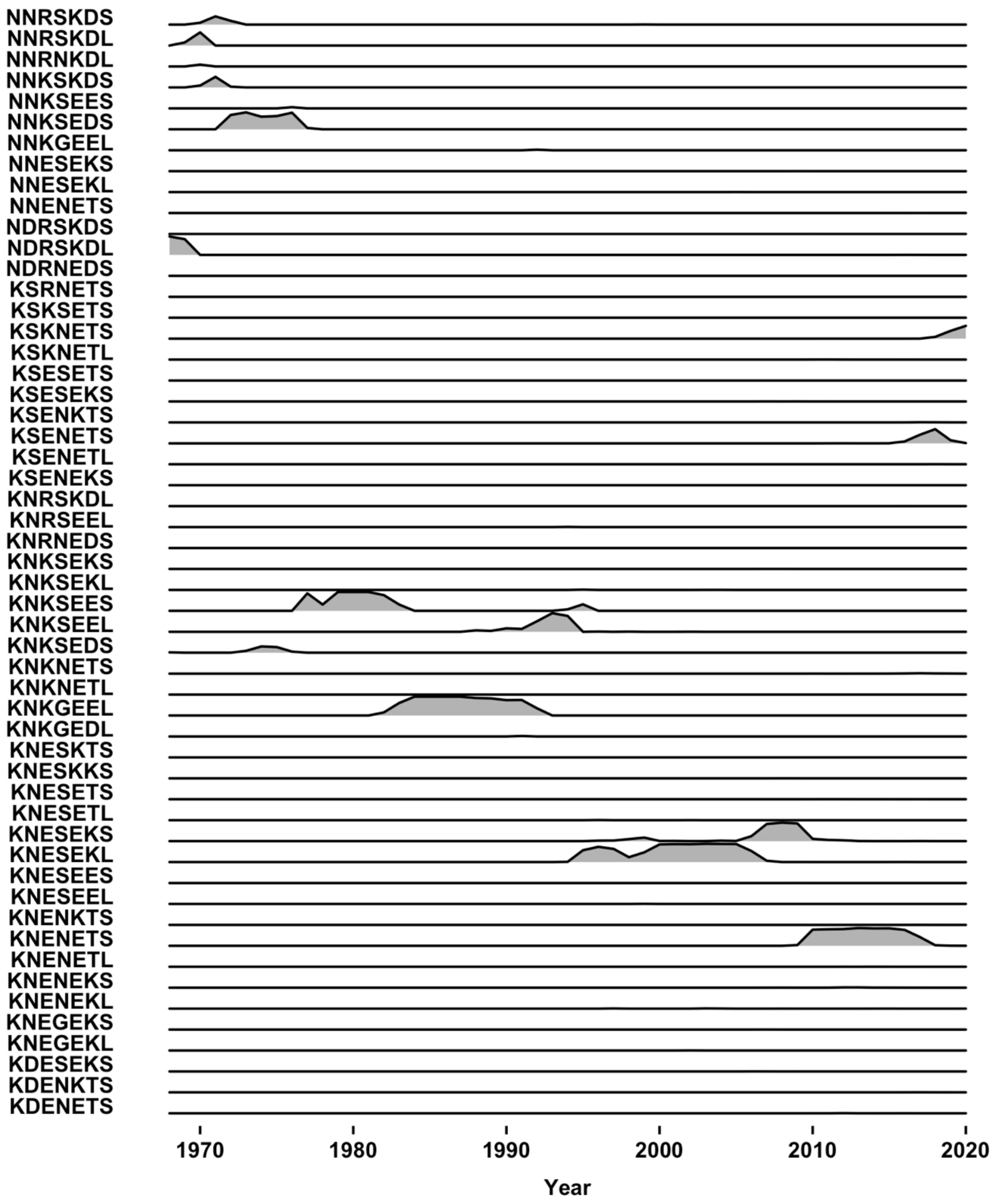


**Supplementary Figure 2. The frequency of haplotypes at the NA antigenic region of interest in circulating human H3N2 strains.** In naturally circulating human H3N2 strains, 53 of the 864 haplotypes of interest were observed. The sequences of these 53 haplotypes are shown on the y-axis. Their occurrence frequency from 1968 to 2020 is plotted. For a frequency of 100%, the density curve would almost touch the next higher baseline. Some haplotypes have a very low occurrence frequency and may look like a complete flat line in the frequency diagram.

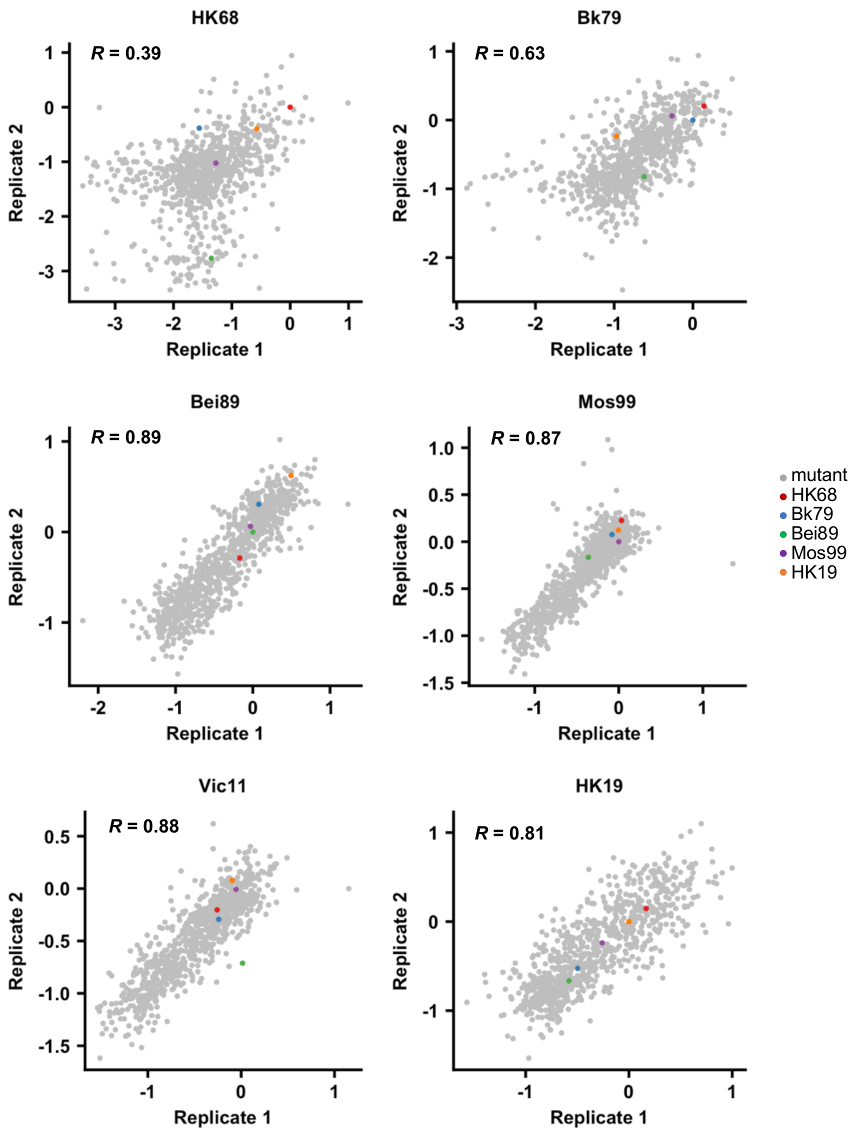


**Supplementary Figure 3. Correlations of fitness measurements between replicates.** Correlations of variant fitness measurements between replicates of deep mutational scanning experiments are shown as scatterplots. Each data point within a scatterplot represents a unique variant. The Pearson correlation (R) between replicates is indicated. Data points corresponding to the WT sequences of HK68, Bk79, Bei89, Mos99, and HK19 are colored as indicated. Of note, the WT sequence of Vic11 contains a naturally rare variant T329 that was not included in our study. Therefore, the WT sequence of Vic11 is not included in our mutant libraries.

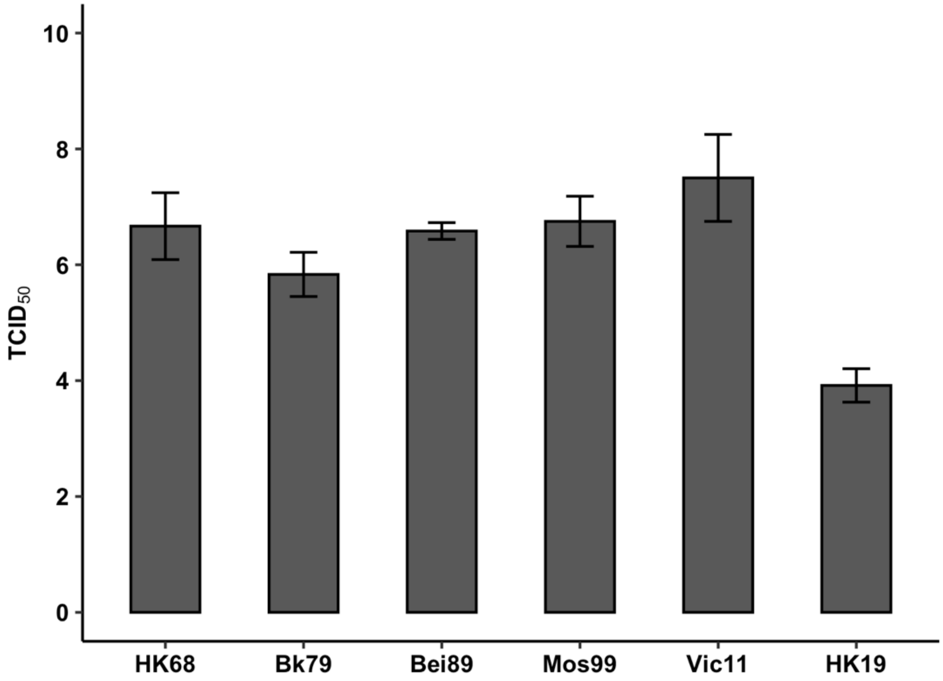


**Supplementary Figure 4. Virus rescue experiment of WT strains.** The titer of six WT strains (HK68, Bk79, Bei89, Mos99, Vic11, and HK19) from a virus rescue experiment is shown. Error bar represents standard error from three biological replicates. This rescue experiment was performed using the six internal segments from A/WSN/33 (H1N1) and HA segment from HK68,

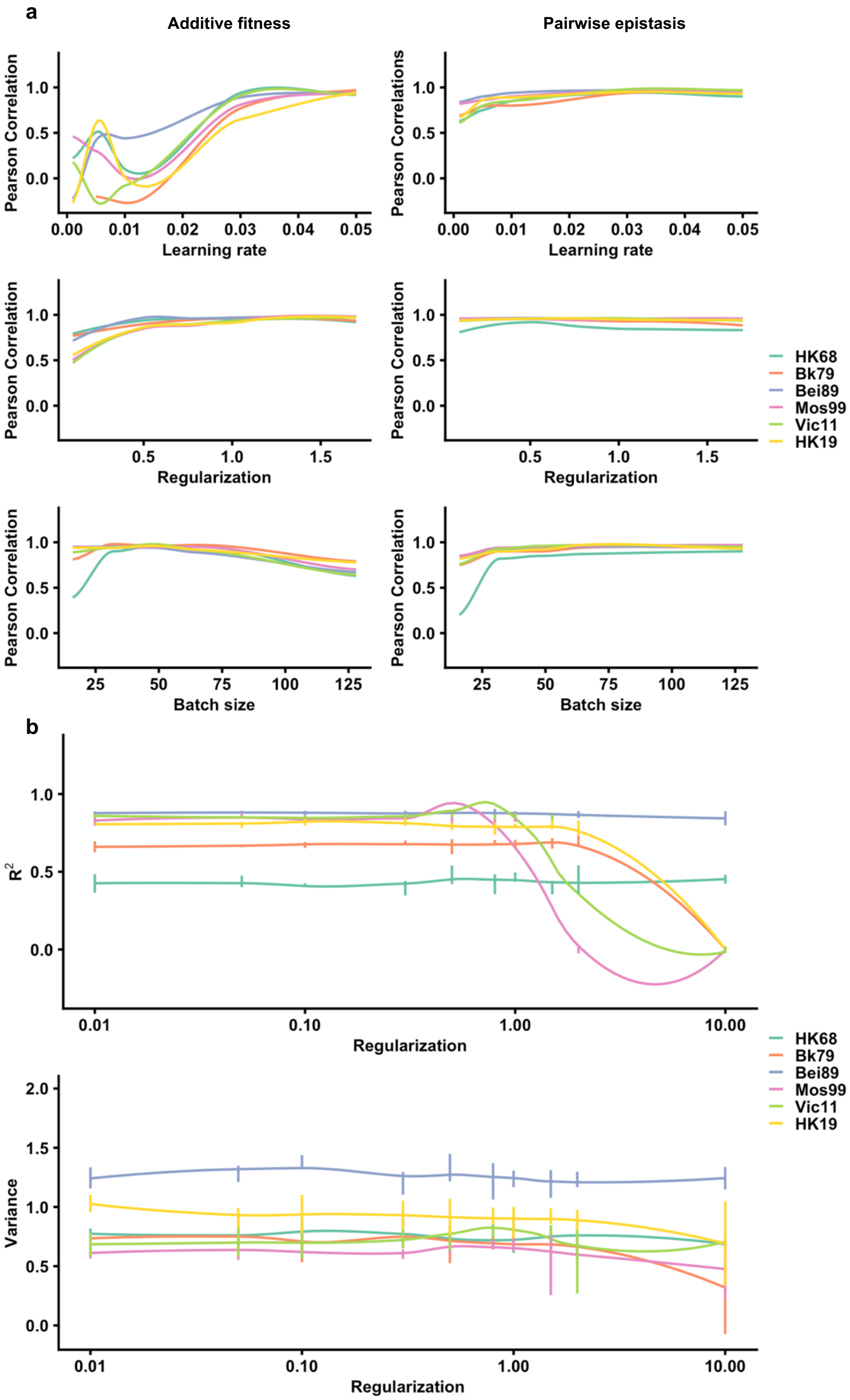


**Supplementary Figure 5. Evaluation of model hyperparameters using repeated k-fold cross-validation. (a)** The relationship between different hyperparameters (learning rate, regularization, and batch size) and model robustness is evaluated. **(b)** The relationship between different regularization hyperparameter and model fit is evaluated.

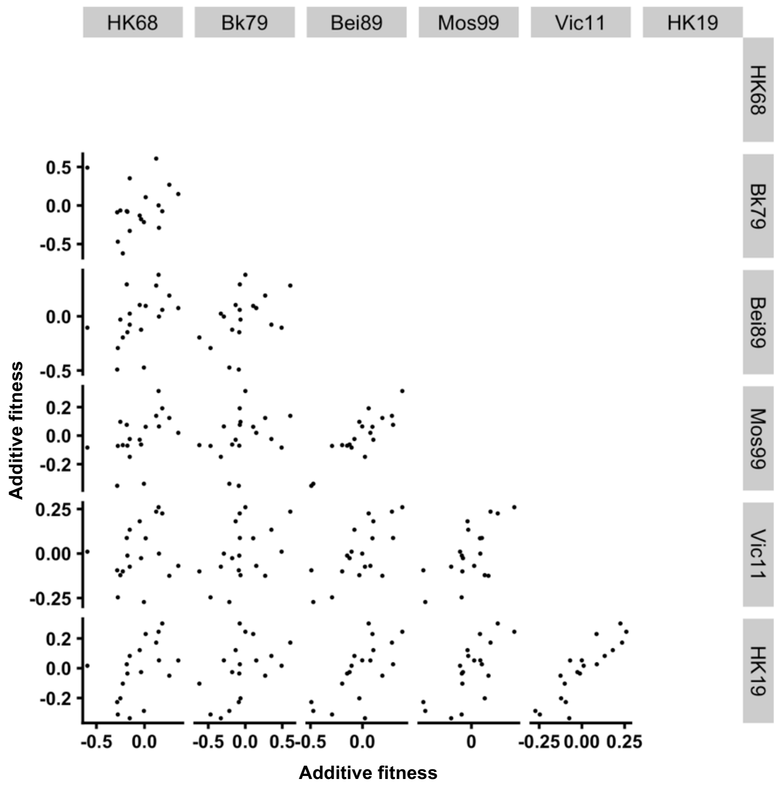


**Supplementary Figure 6. Correlations of additive fitness among genetic backgrounds.** The 19 parameters for additive fitness are compared among genetic backgrounds, with each data point representing one parameter. The correlations of additive fitness among genetic backgrounds varied hugely (Pearson correlation = 0.12 to 0.88).

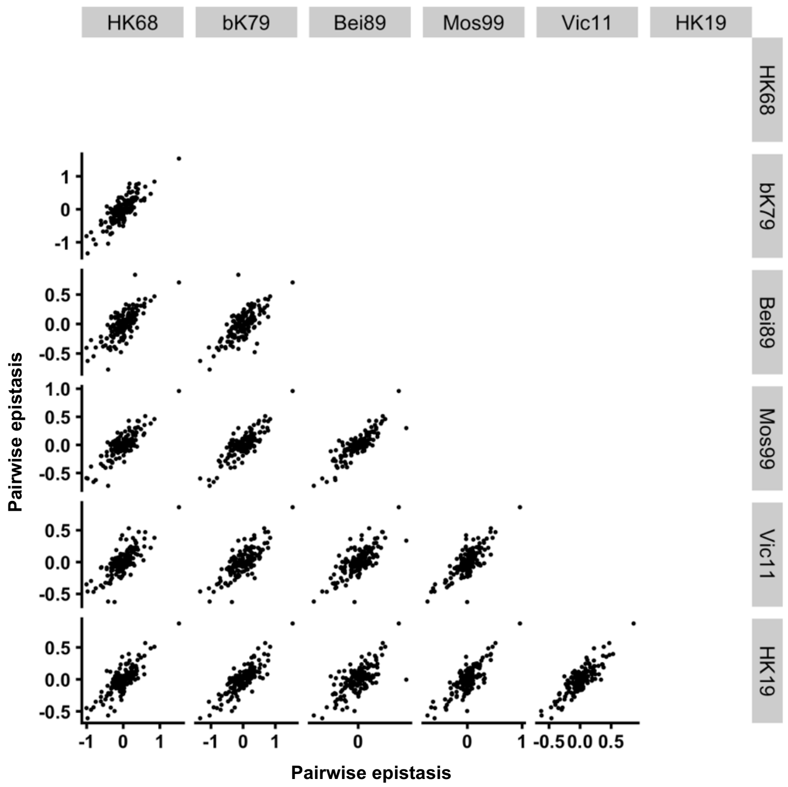


**Supplementary Figure 7. Correlations of pairwise epistasis among genetic backgrounds.** The 153 parameters for pairwise epistasis are compared among genetic backgrounds, with each data point representing one parameter. The correlations of pairwise epistasis among genetic backgrounds are generally strong (Pearson correlation = 0.69 to 0.86).

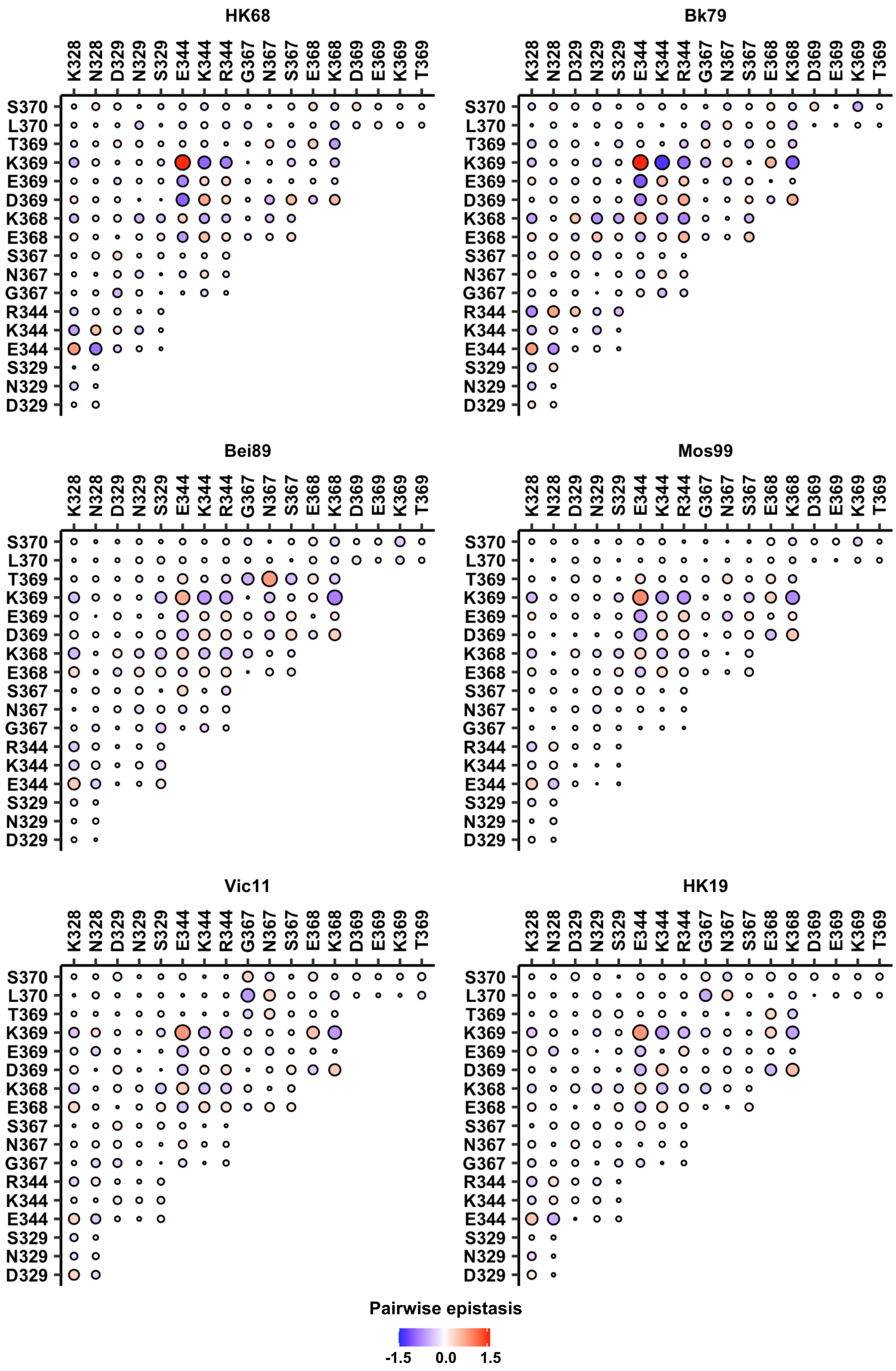


**Supplementary Figure 8. Pairwise epistasis in different genetic backgrounds.** Positive epistasis is in red, while negative epistasis is in blue. The magnitude of epistasis is proportional to size of the circle. The identity of a given double amino acid variant is represented by the labels on the x- and y-axes.

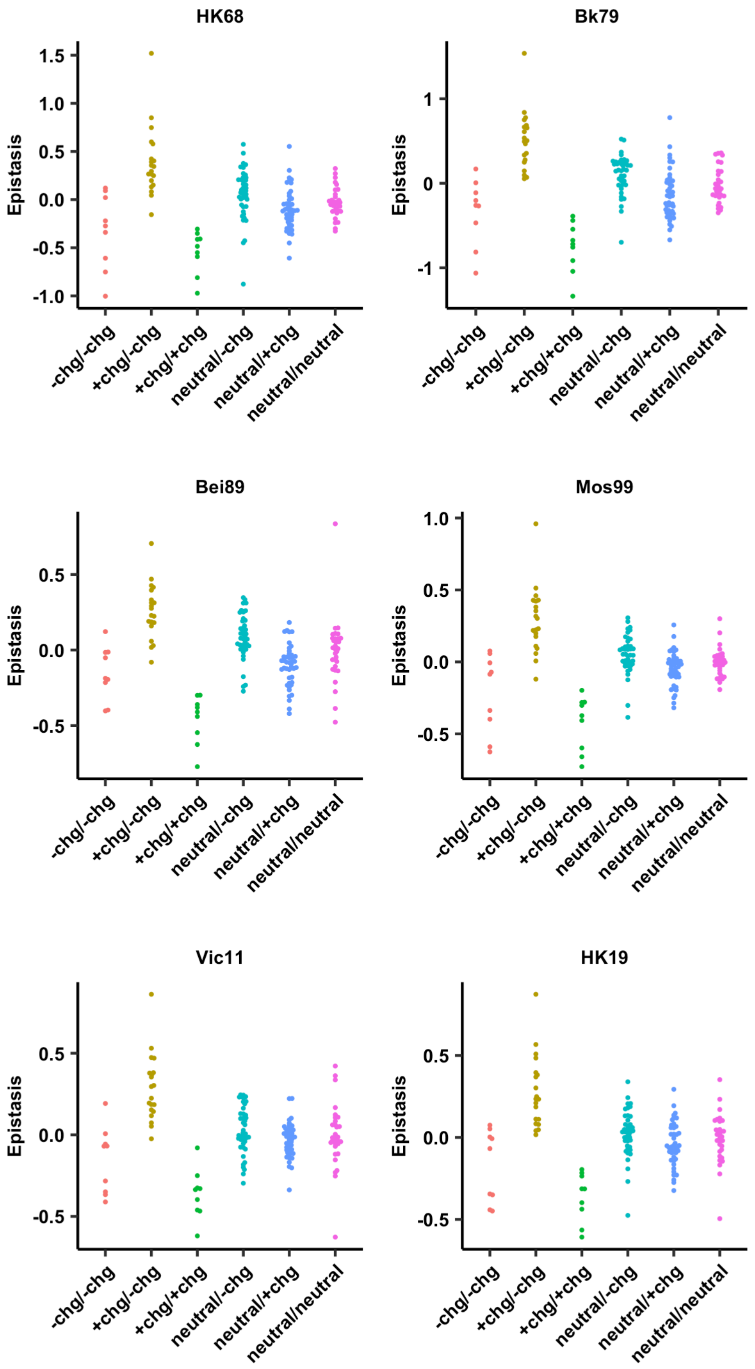


**Supplementary Figure 9. Pairwise epistasis was classified based on the side-chain charge in each pair of amino acid variants.** +chg represents positively charged amino acids (K/R), -chg represents negatively charged amino acids (D/E) and neutral charge represents the remaining amino acids.

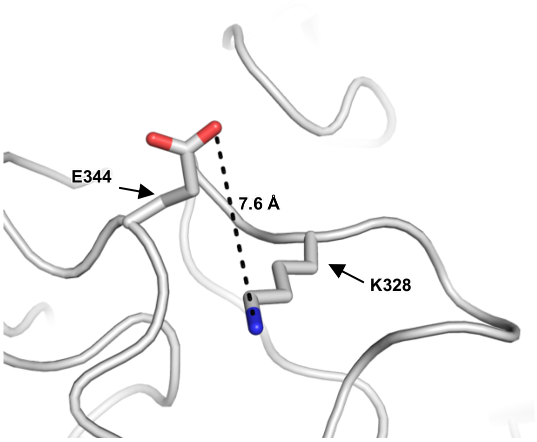


**Supplementary Figure 10.** NA structure from H3N2 A/Tanzania/205/2010 (PDB 4GZO)[1], which has K328 and E344, is shown. The distance between the side chain carboxylate oxygen of E344 and the side chain amine nitrogen of K328 is indicated.

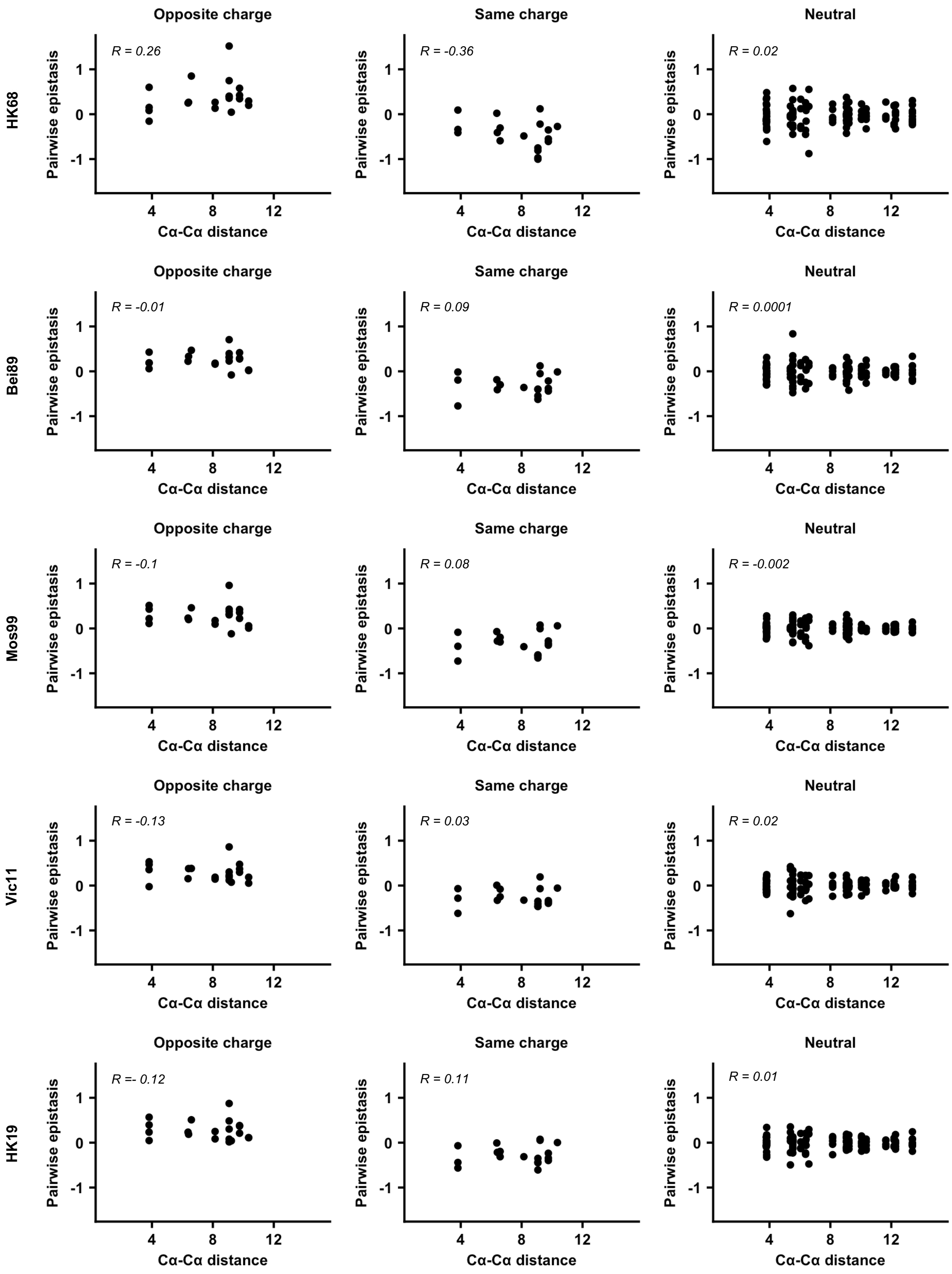


**Supplementary Figure 11. Relationship between the Cα-Cα distance and pairwise epistasis.** The Cα-Cα distance does not correlate with the magnitude of pairwise epistasis, regardless of the genetic background.

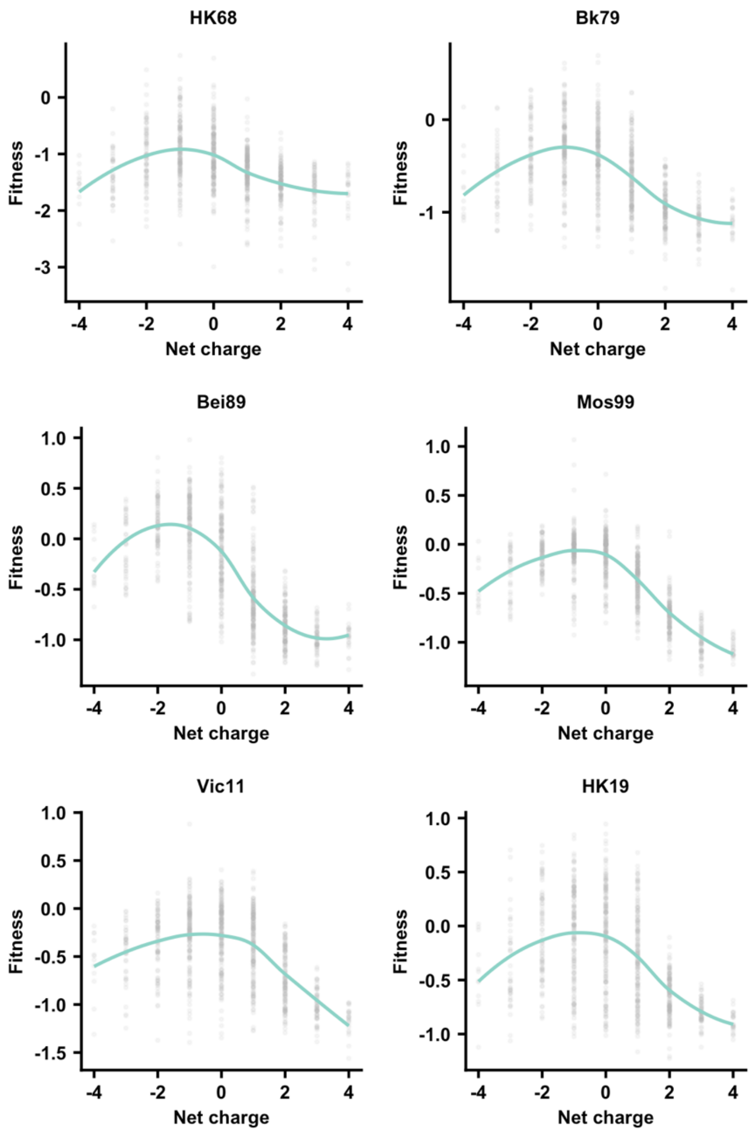


**Supplementary Figure 12. Relationship between variant fitness and the local net charge.** The relationship between variant fitness and the local net charge in the given genetic background is shown. A smooth curve was fitted by loess and shown in teal.

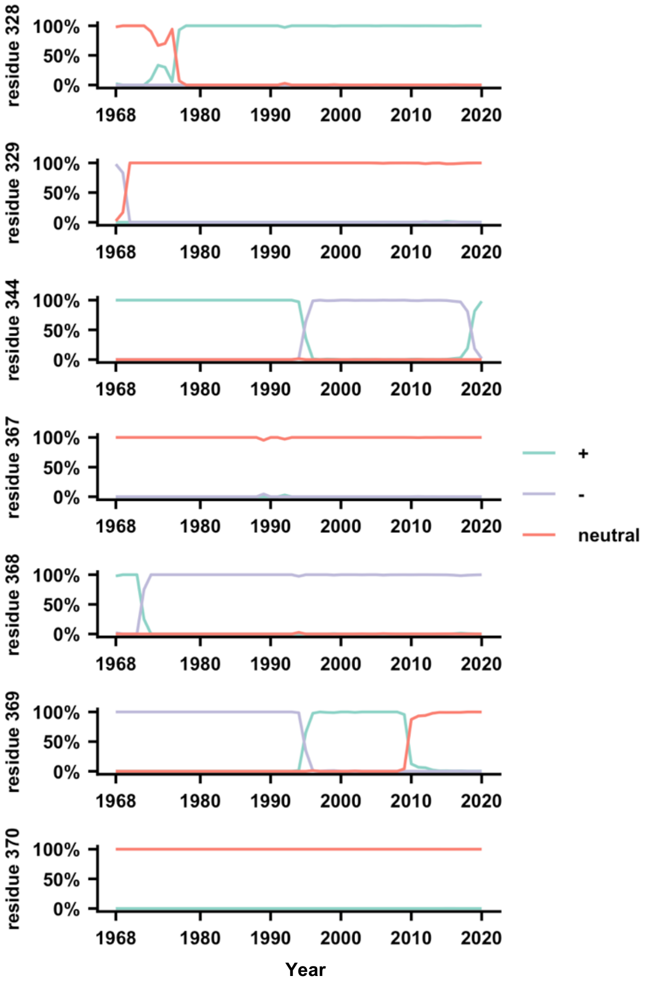


**Supplementary Figure 13. The natural occurrence frequencies of the major amino acid charge states are shown.** (+) represents positively charged amino acids (K/R), (-) represents negatively charged amino acids (D/E), and (n) represents the remaining amino acids.

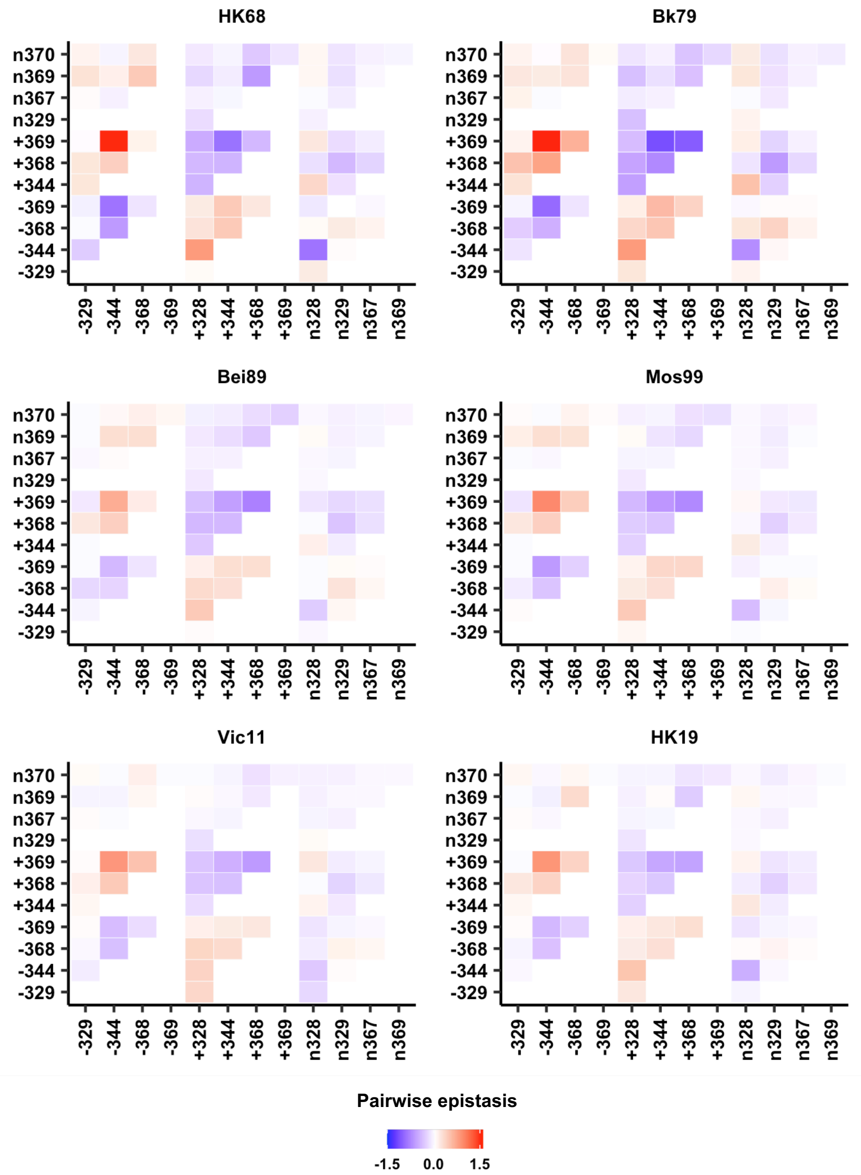


**Supplementary Figure 14. Pairwise epistasis of charged states among seven residues.** Amino acids were classified based on their charge state. (+) represents positively charged amino acids (K/R), (-) represents negatively charged amino acids (D/E), and (n) represents the remaining amino acids. For instance, -344 represents negatively charged amino acids (D/E) at residue 344. The epistasis of +344/-369 would then be the average among that of K344/D369, R344/D369, K344/E369, and R344/E369.

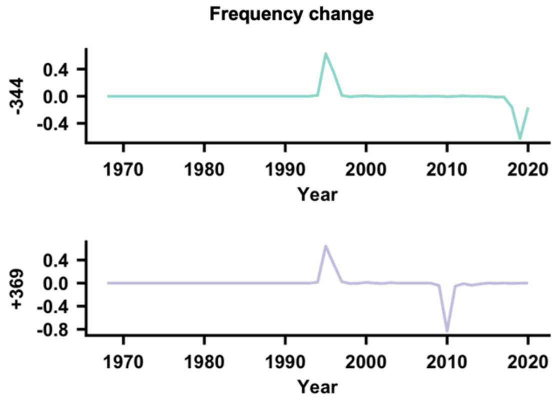


| -344 | +369 | Peak distance (year) | Frequency change |
|---|---|---|---|
| Peak1 | Peak1 | 0 | ++ |
| Peak1 | Peak2 | 15 | +- |
| Peak2 | Peak1 | 24 | -+ |
| Peak2 | Peak2 | 9 | -- |

$$E(-344, +369) = C\sum_{k}^{N} e^{-d_k} = e^{-0} + (-1)e^{-15} + (-1)e^{-24} + e^{-9} = 1.00012$$

**Supplementary Figure 15. Schematic overview of calculating the coevolution score.** The natural frequency changes of amino acids charges at residues 328, 329, 344, 367, 368, 369 and 370 from year to year were calculated. In this simplified example here, one pair of charge states (-344/+369) was used for demonstration. -344 represents negatively charged amino acids (D/E) at residue 344 and +369 means positively charged amino acids (K/R) at residue 369. The frequency change represents frequency in year n minus frequency in year n-1. Both -344 and +369 have two peaks that are above our cutoff of 5% frequency change. Subsequently, peaks from different residues were compared and the distance between each pair of peaks was calculated. Each pair of peaks was classified based on their signs. For example, '++' represents both peaks are positive in magnitude. Eventually, coevolution score of a given pair of charge states was calculated as the sum of exponential decay function of all pairwise peak distances.

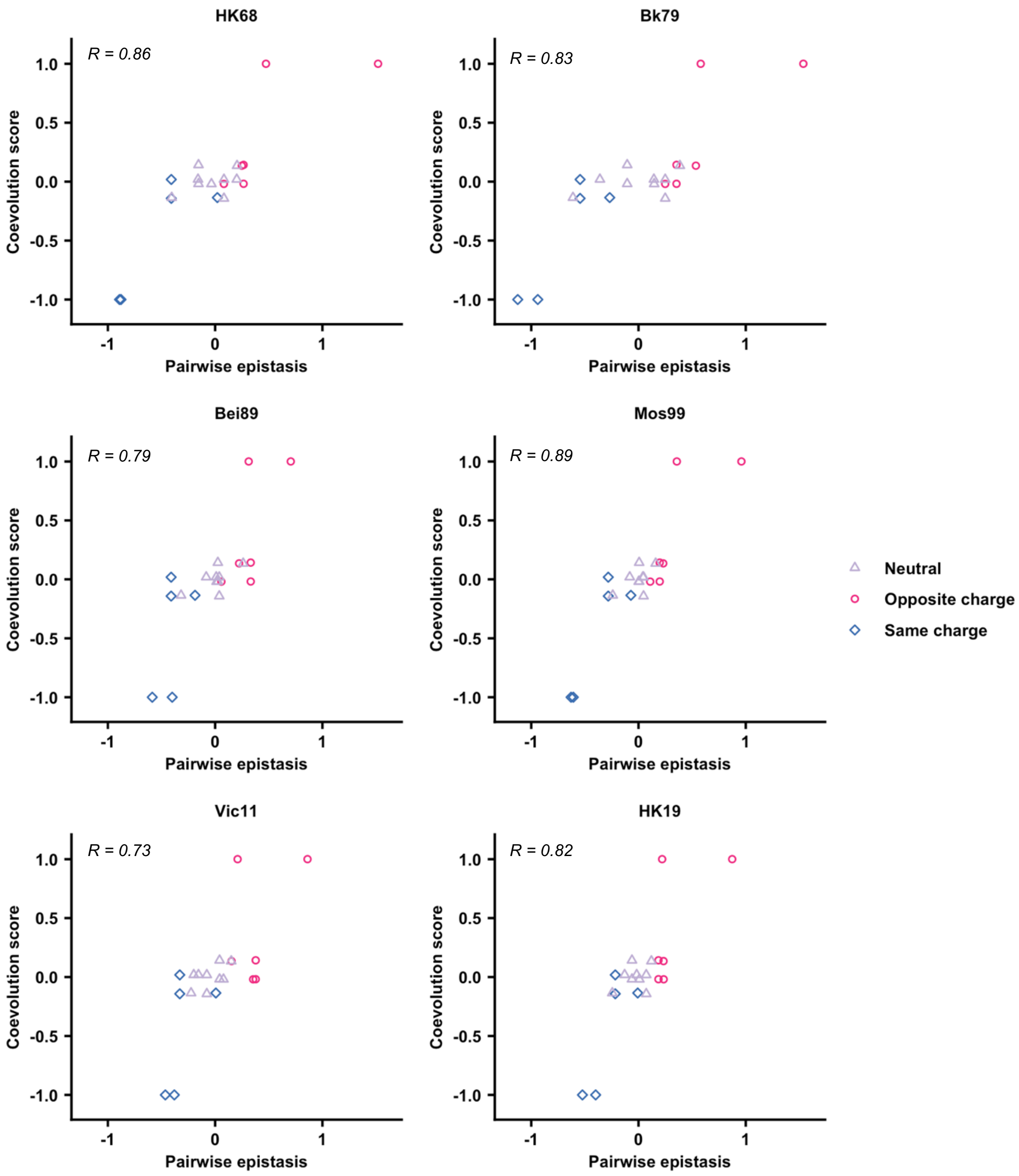


**Supplementary Figure 16. Correlations between coevolution score and pairwise epistasis in different genetic backgrounds.** The relationship between the coevolution score and pairwise epistasis of charge states in the given genetic background is shown. Pearson correlation is indicated.
